# Which cues are sexy? The evolution of mate preference in sympatric species reveals the contrasted effect of adaptation and reproductive interference

**DOI:** 10.1101/2022.10.26.513844

**Authors:** Ludovic Maisonneuve, Charline Smadi, Violaine Llaurens

## Abstract

Mate preferences may target traits (1) enhancing offspring adaptation and (2) reducing heterospecific matings. Because similar selective pressures are acting on traits shared by different sympatric species, preference enhancing offspring adaptation may increase heterospecific mating, in sharp contrast with the classical case of so-called ‘magic traits’. Using a mathematical model, we study which and how many traits will be used during mate choice, when preferences for locally adapted traits increase heterospecific mating. In particular, we study the evolution of preference towards an adaptive *vs*. a neutral trait in sympatric species. We take into account sensory trade-offs which may limit the emergence of preference for several traits. Our model highlights that the evolution of preference towards adaptive *vs*. neutral traits depends on the selective regimes acting on traits but also on heterospecific interactions. When the costs of heterospecific interactions are high, mate preference is likely to target neutral traits that become a reliable cue limiting heterospecific matings. We show that the evolution of preference towards a neutral trait benefits from a positive feedback loop: the more preference targets the neutral trait, the more it becomes a reliable cue for species recognition. We then reveal the key role of sensory trade-offs and the cost of choosiness favouring the evolution of preferences targeting adaptive traits, rather than traits reducing heterospecific mating. When sensory trade-offs and the cost of choosiness are low, we also show that preferences targeting multiple traits evolve, improving offspring fitness by both transmitting adapted alleles and reducing heterospecific mating. Altogether, our model aims at reconciling ‘good gene’ and reinforcement models to provide general predictions on the evolution of mate preferences within natural communities.

**Impact Summary:** Mate preferences are widespread throughout the animal kingdom and generate powerful selective forces impacting the diversification of traits and species. The evolution of such preferences has been the focus of multiple theoretical and empirical studies and intense scientific debates. The evolution of mate preference (1) enhancing offspring fitness and (2) reducing heterospecific mating have been mostly studied separately, except in the specific case of preference for so-called ‘magic traits’ that increase both offspring survival and species divergence. However, in many cases, the evolution of traits in sympatric species generates conflicting evolutionary forces acting on preferences. On one hand, enhanced offspring survival promotes preference towards locally adaptive traits and may thus lead to convergent evolution of traits among sympatric species. On the other hand, the evolution of similar traits in sympatric species may generate costly heterospecific sexual interactions promoting preference towards traits that diverge between species. Here, we thus build a general mathematical model to investigate the evolutionary factors determining which and how many traits are targeted by mate choice. We especially determine whether preferences will likely target adaptive *vs*. neutral traits. Our model highlights that the evolution of preferences for adaptive *vs*. neutral traits in sympatric species depends on within-species mating opportunities but also on the niche overlap between species, tuning heterospecific interactions. By jointly considering (1) the selection regimes acting on the targeted traits within species, as well as (2) interactions with other species living in sympatry, our theoretical study provides a general framework reconciling these research fields.

## Introduction

The evolution of mate preference plays a major role in diversifying traits (Chouteau et al., 2017) and species (Stre et al., 1997). Nevertheless, we still know little about the evolutionary factors determining the traits preferentially targeted by preferences. Preferences target traits displayed by the parents, but their evolution may depend on the indirect fitness benefit brought to the offspring carrying locally adapted traits (Neff and Pitcher, 2005). The evolution of mate preference depends not only on intraspecific competition but also on the ecological interactions with sympatric species. When species occur in sympatry, sexual interactions with heterospecifics (Gröning and Hochkirch, 2008) can lead to fitness reduction because of limited survival in the resulting hybrids (Merrill et al., 2012), but also because of costly heterospecific courtship and rivalry. These fitness costs generated by heterospecific interactions can promote mate preferences targeting traits that differ between species (McPeek and Gavrilets, 2006; Yamaguchi and Iwasa, 2013). The evolution of preferences may thus depend on both (1) the selection regimes acting on the targeted traits within species and (2) the distribution of these traits in other species living in sympatry. Such multifactorial selection acting on the different traits displayed by males may then favour the evolution of female preferences targeting several traits. Preference for multiple cues may improve some components of fitness in the offspring and/or limit heterospecific matings (Candolin, 2003).

However, by contrast with classical ‘magic’ traits under disruptive selection (Servedio et al., 2011), natural selection frequently promotes similar traits in different sympatric species (*e*.*g*. in mimetic species, (Sherratt, 2008)). Indirect fitness benefits enjoyed by the offspring may promote preference increasing the risk of heterospecific matings (*e*.*g*. (Gumm and Gabor, 2005; Higgie and Blows, 2007)). Preferences targeting multiple traits may then improve offspring fitness by both transmitting adapted alleles and reducing heterospecific mating (Candolin, 2003). However, several constraints might limit the number of traits targeted by preference (Schluter and Price, 1993; Pomiankowski and Iwasa, 1993; Iwasa and Pomiankowski, 1994). The number of available partners displaying the preferred combination of traits may also impact the evolution of preference for multiple traits. The cost of choosiness associated with rejecting unpreferred males by choosy females may increase when the number of targeted traits grows. The complex cognitive processes involved may also limit the evolution of preference for multiple traits (Crapon de Caprona and Ryan, 1990).

Here, we propose a general mathematical framework to predict the trait targeted by mate choice, when preferences for locally adapted traits increase heterospecific mating. We use mathematical modelling to investigate the evolution of preference based on multiple traits, assuming sensory trade-offs impacting the relative perception of different mating cues. We study the evolution of preference towards two evolving traits (*T*_1_ and *T*_2_) shared by a pair sympatric species (*A* and *B*). We assume that selection regimes acting on traits increase species similarity. We aim at identifying how selection regimes acting on the targeted traits, as well as reproductive interference between species, favour (1) preference for locally adapted trait *vs*. trait reducing heterospecific mating and (2) preference for single *vs*. multiple traits.

## Method

### Model overview

We consider two sympatric species (*A* and *B*) and assume that individuals from both species display two traits that female preferences could target. We assume that both traits are controlled by independent haploid loci (called *T*_1_ and *T*_2_, respectively), with two possible alleles 0 or 1 at each locus. For the sake of simplicity, we assume that alleles 0 or 1 code for trait values 0 or 1, respectively. We fix the genotypic distribution in species *B* by assuming that all individuals carry the allele 1 at both trait loci. We assume that a parameter *ρ* modulates the strength of female preferences. The loci *P*_1_ and *P*_2_ then determine the value of traits *T*_1_ and *T*_2_, respectively, preferred by females. Female preference can target either or both traits (*T*_1_ and *T*_2_) displayed by the males: a preference locus *M* controls the relative level of attention paid by the female toward trait *T*_1_ *vs*. trait *T*_2_ expressed by males. We thus introduced *γ* as the *relative preference weighting*, modulating the level of attention on either trait (Figure 1). We assume that a resident and a mutant alleles can occur at locus *M* with different values of *γ* (*γ*_r_ and *γ*_m_). We then study the evolution of the *relative preference weighting* in focal species *A*. Assuming non-overlapping generations and relying on an adaptive dynamics framework, we investigate the fixation of new mutant alleles and estimate the equilibrium *relative preference weighting* (*γ*^∗^). The ancestral *relative preference weighting* is given by 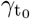.

**Figure 1.**
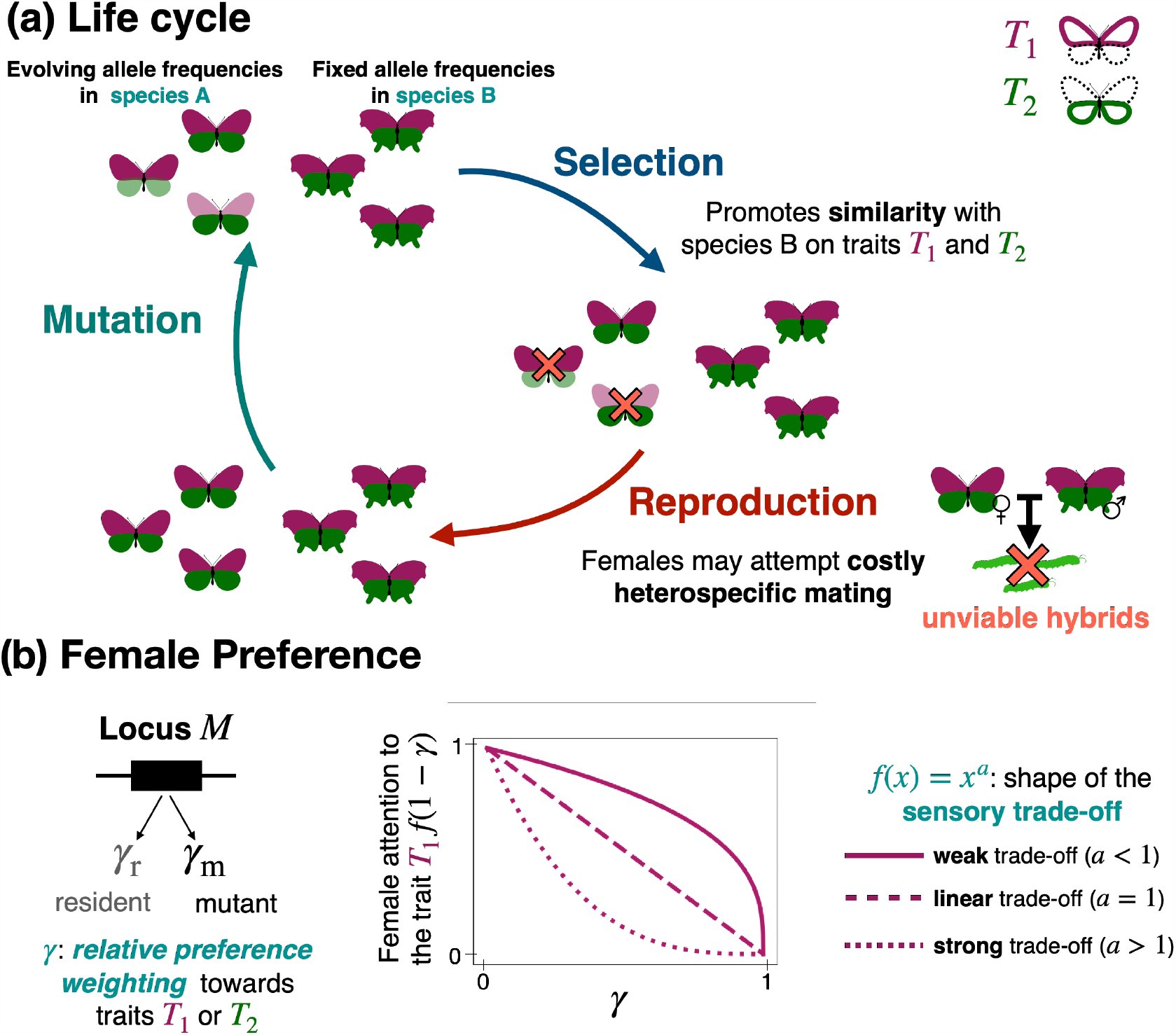
Schematic description of the model. (a) Life cycle. The evolution of the preference may depend on the interactions between species and natural selection acting on the preferred traits. Here, we assume two sympatric species, *A* and *B*, depicted in this scheme as butterfly species with different wing shapes. The individuals can display two traits, *T*_1_ and *T*_2_, represented by the forewing and hindwing colours as an example. We study the coevolution of the trait values (0 or 1, shown as intense *vs*. light colour of the wings) and the preference in species *A*. We assume that all individuals in species *B* displayed the trait value 1 (intense colour) at both traits. We assume a selection step, promoting trait value 1 at both traits in species *A*, increasing similarity with species *B*, where value 1 is fixed for both traits. We then assume a reproduction step, where the mating success of the different individuals in species *A* depends on the traits and preferences carried by males and females. In particular, females of species *A* may attempt costly and unfertile sexual interactions with males of species *B* depending on their preferences. (b) Genetic basis of preference. Depending on the genotype at locus *M*, females modulate their level of attention towards either trait displayed by males (*relative preference weighting γ*). We also assume that the level of attention on one trait diminishes the attention on the alternative one. We investigate several shapes of this trade-off tuned by the parameter *a*.

The evolution of the *relative preference weighting* depends on the survival of the produced offspring. We thus assume that the traits *T*_1_ and *T*_2_ displayed by the individuals can modify their survival. *s*_1_ and *s*_2_ gives the selective advantages of allele 1 at locus *T*_1_ and *T*_2_, respectively. We assume that *s*_1_ ≥0 and *s*_2_ ≥0 so that natural selection acts within species *A* promotes similarity with species *B* (recall that allele 1 is fixed for both traits in species *B*). We also assume that females can encounter and have sexual interactions with heterospecifics. Such sexual interactions lead to fitness costs but do not produce any viable offspring.

To model a sensory trade-off acting on preferences, we also assume that the level of attention on one trait diminishes the attention on the alternative one. We investigate several shapes of this trade-off tuned by the parameter *a* (Figure 1). Finally, we assume that after refusing a mating opportunity females may not encounter another male with a probability *c*, leading to a cost of choosiness. We investigate how the cognitive trade-off and the costs of choosiness impact the evolution of the direction of preference, which might lead to a preference for either a single or multiple traits.

### Selection regime acting on the displayed traits

We assume that individuals display two different traits *T*_1_ and *T*_2_, each controlled by a single biallelic locus. We assume that the traits *T*_1_ and *T*_2_ displayed by the individuals can modify their survival. Let *𝒢* = {0, 1}^5^, be the set of all genotypes at loci *T*_1_, *T*_2_, *P*_1_, *P*_2_ and *M*. We define *f*_*i*_ and 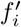 as the frequencies of genotype *i* ∈ *𝒢* in the focal species before and after a step of natural selection acting on survival, respectively. The resulting frequency after selection, 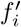 is then given by

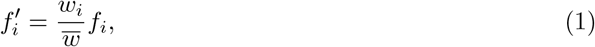

where *w*_*i*_ is the fitness of an individual of genotype *i* during natural selection, and the mean fitness 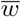 is a normalisation term to ensure that genotype frequencies sum to 1 after selection.

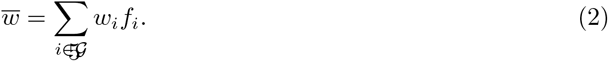

We recall that *s*_1_ and *s*_2_ are the selective advantages of allele 1 at locus *T*_1_ and *T*_2_, respectively. The fitness component due to the natural selection of an individual of genotype *i* is thus given by:

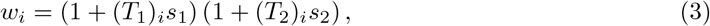

where (*T*_1_)_*i*_ and (*T*_2_)_*i*_ refer to the allele (0 or 1) an individual of genotype *i* carries at loci *T*_1_ and *T*_2_ respectively (following Kirkpatrick et al. (2002)). For example, (*T*_1_)_*i*_ = 1 and (*T*_2_)_*i*_ = 0 for an individual carrying allele 1 at locus *T*_1_ and allele 0 at locus *T*_2_.

### Reproductive success depending on female preference on traits displayed by males

#### Preference based on multiple traits

Females generally use both traits in mate choice but may vary in their relative attention given to trait *T*_1_ *vs*. trait *T*_2_. This relative attention depends on the *relative preference weighting* parameter *γ*. Alleles at locus *M* determine the *relative preference weighting* : allele 0 (resp. 1) is associated with the value *γ*_r_ (resp. *γ*_m_). Translating into an equation, the *relative preference weighting* of a female of genotype *j* is thus given by:

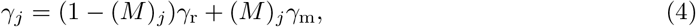

where (*M*)_*j*_ is the allele (0 or 1) at locus *M* in genotype *j*. We assume that a cognitive trade-off, described by the function *f*, also impacts the relative attention to the two traits. *f* (1 −*γ*) and *f* (*γ*) determine the attention on trait *T*_1_ and *T*_2_ where *a* ∈ [0, +∞) and

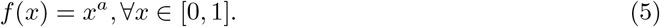

The function *f* is non-decreasing, so attention on one trait diminishes attention on the alternative trait. Moreover, *f* (0) = 0 and *f* (1) = 1, so the female choice relies on a single trait in the two extreme cases. The parameter *a* tunes the shape of the trade-off function *f* (Figure 1):

- when *a* = 1, *f* is linear, leading to a **linear trade-off**, where the female attention on traits 1 (resp. 2) is proportional to 1 − *γ* (resp. *γ*).
- when *a <* 1, *f* is concave, leading to a **weak trade-off** between attention towards the two male traits. Females can thus use both traits for mate choice.
- when *a >* 1, *f* is convex, leading to a **strong trade-off** in female attention between the two traits. Females focusing on one trait largely ignore the alternative trait, and intermediate values of *γ* lead on poor attention to both traits.

Therefore, when a female of genotype *j* encounters a male of genotype *k*, she accepts the male with probability

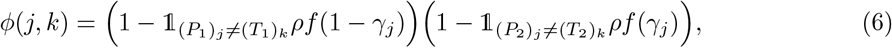

where 𝟙_{.}_ is the indicator function that returns 1 if the condition in the subscript is realised and 0 otherwise. The mating probability for a pair composed of a female with genotype *j* and a male with genotype *k* depends on the match between the female’s preferred traits (given by (*P*_1_)_*j*_ and (*P*_2_)_*j*_) and the male’s traits values (given by (*T*_1_)_*k*_ and (*T*_2_)_*k*_). When the female does not prefer the male traits (either (*P*_1_)_*j*_ ≠ (*T*_1_)_*k*_ or (*P*_2_)_*j*_ ≠ (*T*_2_)_*k*_), the female tends to reject the male. The parameter *ρ* quantifies the strength of female preference.

#### Mating process

We assume that females can mate at most once. We assume that each female sequentially meets males in a random order. These males can be either conspecifics or heterospecifics (see Figure 2 for an illustration of the mating process). At each encountering event, the female may accept the male with a probability depending on her preference and on the traits displayed by the encountered male.

**Figure 2.**
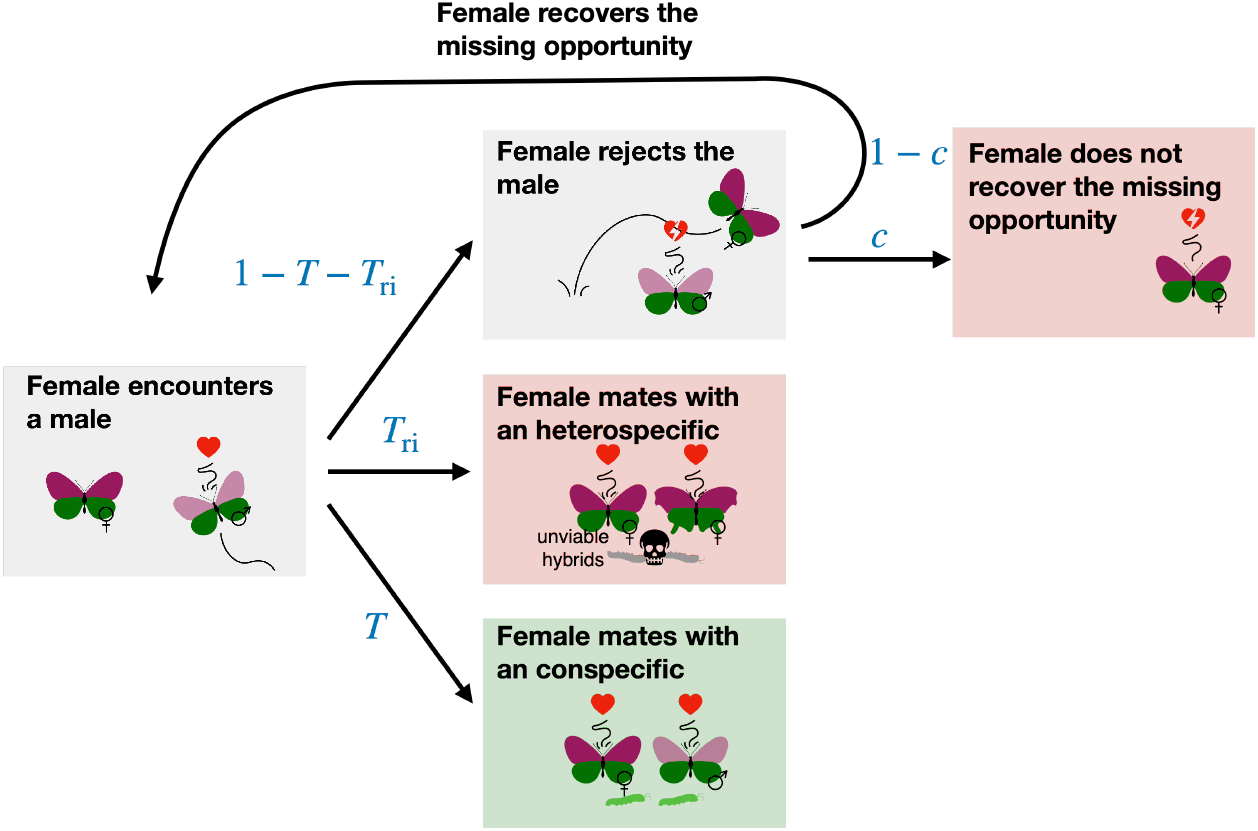
Illustration of the mating process. We assume that females can mate at most once. We assume that each female meets sequentially random males, which can be either conspecifics or heterospecifics. At each encounter, the female accepts or rejects the male with a probability depending on the female’s preference and the male’s traits. After an encounter, a female accepts a conspecific (resp. heterospecific) male with probability *T* (resp. *T*_ri_). If a female engages in heterospecific mating, she produces totally unviable offspring. Because females mate at most once, a female engaged in a heterospecific mating cannot recover the associated fitness loss. Moreover, we assume that females refusing a mating opportunity can encounter another male with a probability of 1 − *c*. We interpret *c* as the cost of choosiness.

Given that a female of genotype *j* encounters a male uniformly at random, *T* (*j*) gives the realized probability that a female of genotype *j* accepts a conspecific (Otto et al., 2008):

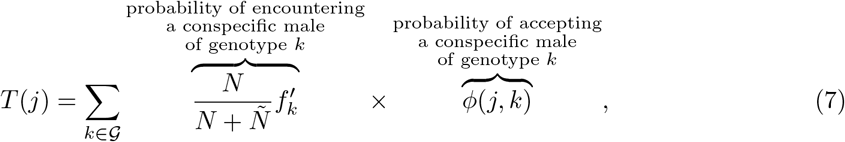

where *N* and *Ñ* are species *A* and *B* densities, respectively. *T* (*j*) depends on (1) the probability of encountering a conspecific *vs*. a heterospecific male, (2) on the distribution of traits within conspecific males and (3) on female preference.

A female of genotype *j* may also accept a heterospecific male with a probability

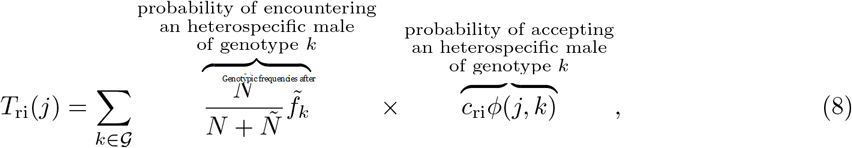

where *c*_ri_ ∈ [0, 1] modulates the probability for the female to accept heterospecific males. This parameter may capture the effect of other unmodelled traits the females can assess, and tunes the strength of reproductive interference. We assume a same genetic architecture of traits in species *A* and *B*, 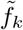 is the frequency of genotype *k* in species *B*. We assume that the allele 1 is fixed at both trait loci (*T*_1_ and *T*_2_) in species *B*. We assume that heterospecific crosses produce totally unviable offspring. Because females mate at most once, a female engaged in a heterospecific mating cannot recover the associated fitness loss.

However, we assume that females refusing a mating opportunity encounter another male with a probability of 1 − *c*. We interpret *c* as the cost of choosiness. The probability that a female of genotype *j* mates with a conspecific male is thus given by

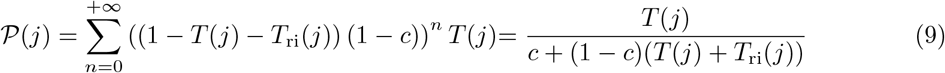

where ((1 − *T* (*j*) − *T*_ri_(*j*)) (1 − *c*))^*n*^ is the probability that a female of genotype *j* rejects the *n* males she first encounters and then encounters an (*n* + 1) − *th* male.

#### Mating success of a pair

We now compute the mating success of a pair. Knowing that a female of genotype *j* has mated with a conspecific male, the probability that this male is of genotype *k* is given by

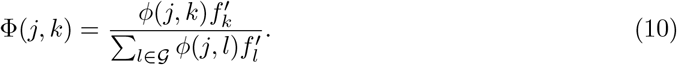

The contribution to the next generation of a mating between a female of genotype *j* and a male of genotype *k m*_*j,k*_ is thus given by the product of (1) the probability that a female of genotype *j* mates with a conspecific male 𝒫 (*j*) with (2) the probability the female mates with a male of genotype *k* knowing that the female has mated with a conspecific male Φ(*j, k*)

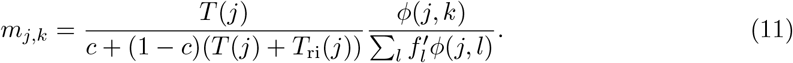

Genotypic frequencies after reproduction in the focal species then depend on the contribution to the next generation of the different crosses between females and males of genotype *j* and *k*, respectively, described by *m*_*j,k*_, for all *j* and *k* in 𝒢. We note 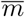 the mean value of this contribution across all mating pairs

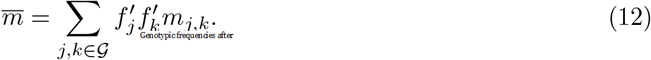

The frequency after reproduction of genotype *i* in species A is then given by

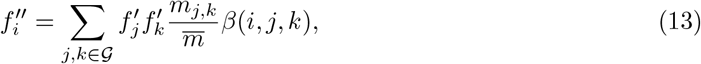

where *β*(*i, j, k*) the probability that a mating between a female of genotype *j* and a male of genotype *k* produces an offspring of genotype *i. β*(*i, j, k*) describes the segregation of alleles during reproduction. We assume recombination between female and male haplotypes. The offspring inherits any of the two recombined haplotypes with a probability one half.

### Mutation

We assume that mutations occur at loci *T*_1_, *T*_2_, *P*_1_ and *P*_2_ within offspring with probability 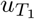, 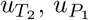 and 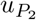 respectively.

We summarise all the model’s variables and parameters in Table A1.

### Model exploration

#### Invasion analyses

In an adaptive dynamics framework, we study the invasion of a rare mutant at locus *M* associated with the value of *relative preference weighting γ*_m_ in a resident population where the resident allele codes for the value *γ*_r_. We assume that the mutation has a small effect, so that *γ*_r_ − *γ*_m_ is small. Before the mutant introduction, we assume that genotypic frequencies at loci *T*_1_, *T*_2_, *P*_1_ and *P*_2_ evolve toward equilibrium allelic frequency values named 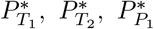 and 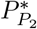. Initially, we assume no genetic association. For the parameters space explored in this study, the initial allele frequencies have no impact on the evolutionary dynamics (see Appendix A2). Studying mutant invasion for all possible resident populations, we estimate the different types of singular *relative preference weightings* numerically (Rousset, 2004; Geritz et al., 1998):

- Continuously stable *relative preference weighting* : The model predicts a convergent evolution toward this gamma value, and once this value is reached, any mutant associated with other gamma value will then fail to invade.
- Evolutionary repeller: The model predicts that the evolution of gamma will always strongly depart from this value.
- Branching point: The model predicts a convergent evolution toward this gamma value, but once this value is reached, mutants associated with other gamma value will invade, leading to polymorphism (never observed in this study).

We determine the equilibrium *relative preference weighting* (*γ*^∗^) *i*.*e*. the continuously stable *relative preference weighting* reached by the dynamics that may depend on ancestral preference 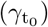. By default, we assume that ancestral preference equally targets both traits 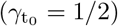.

#### QLE analysis

Assessing mutant invasion would first require to estimate the 2^4^ − 1 = 15 equilibrium genotypic frequencies at loci *T*_1_, *T*_2_, *P*_1_ and *P*_2_ in the resident population (with the resident allele fixed at locus *M*). Then, the 2^5^ − 1 = 31 genotypic frequencies at loci *T*_1_, *T*_2_, *P*_1_, *P*_2_ and *M* would need to be tracked down throughout the generations following mutant introduction, to determine whether the mutant allele invades. To simplify the model analyses, we perform a Quasi-Linkage Equilibrium (QLE) analysis. The QLE approach allows to analytically estimate the change in allele frequencies and in genetic associations (Kirkpatrick et al., 2002). Under the QLE hypotheses, the genetic associations quickly reach their equilibrium value (Nagylaki, 1993), so they can be approximated by their equilibrium value (Kirkpatrick et al., 2002). Using the estimated change of allele frequencies at loci *T*_1_, *T*_2_, *P*_1_ and *P*_2_ under QLE approximation, we compute the equilibrium allele frequencies at these loci before mutant introduction. We then use the sign of the change of mutant allele frequency in the resident population under QLE approximation to assess mutant invasion.

We compute the selection gradient *S* by rewriting the approximated change of mutant frequency under QLE (given in Equation (A10))

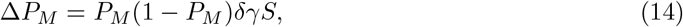

where *δγ* = *γ*_m_ − *γ*_r_ is the effect of the mutant allele on the *relative preference weighting*. When the selection gradient is positive (resp. negative), selection promotes the evolution of preference towards the trait *T*_2_ (resp. *T*_1_). Using the selection gradient *S*, we can disentangle the relative effects of the different selection pressures acting on the evolution of the *relative preference weighting*. We decompose the selection gradient in four terms

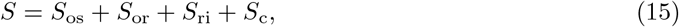

where *S*_os_, *S*_or_, *S*_ri_ and *S*_c_ (those expression are given in Equations (A21), (A22), (A16) and (A18)), captures the effect of offspring survival, offspring reproductive success, reproductive interference and cost of choosiness on mutant fitness respectively. More precisely, *S*_os_ and *S*_or_ capture the indirect fitness benefit of producing offspring carrying locally adapted and sexy traits, respectively. *S*_ri_ and *S*_c_ capture the direct fitness benefit for a female of reducing heterospecific mating and the cost of choosiness.

The QLE analysis assumes that selection is weak and recombination is strong compared to selection. In line with this hypothesis, we assume that *s*_1_, *s*_2_, *ρ, c*_ri_, *c* are of order *ε* with *ε* being small and recombination rates of order 1. We also assume that mutation rates are of order *ε*. We perform the QLE analysis using Wolfram Mathematica 12.0, and provide detailed results of these analyses in Appendix A1.

To check the robustness of specific results, we also run numerical analyses, relaxing the QLE assumptions (*e*.*g*. Figure A5).

#### Default parameters

If not specified we use the following parameter values: 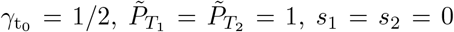, 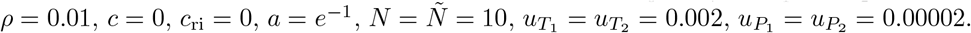.

## Results

### ’Good genes’ *vs*. reinforcement

To explore the evolution of preference enhancing offspring fitness *vs*. reducing heterospecific mating, we computed the equilibrium value of the *relative preference weighting γ*^∗^ when assuming that natural selection acts on trait *T*_1_, increasing similarity with species *B* (*s*_1_ *>* 0), while the trait *T*_2_ is neutral (*s*_2_ = 0). We investigate different strengths of natural selection acting on trait *T*_1_ (*s*_1_) and reproductive interference (*c*_ri_) (Figure 3).

**Figure 3.**
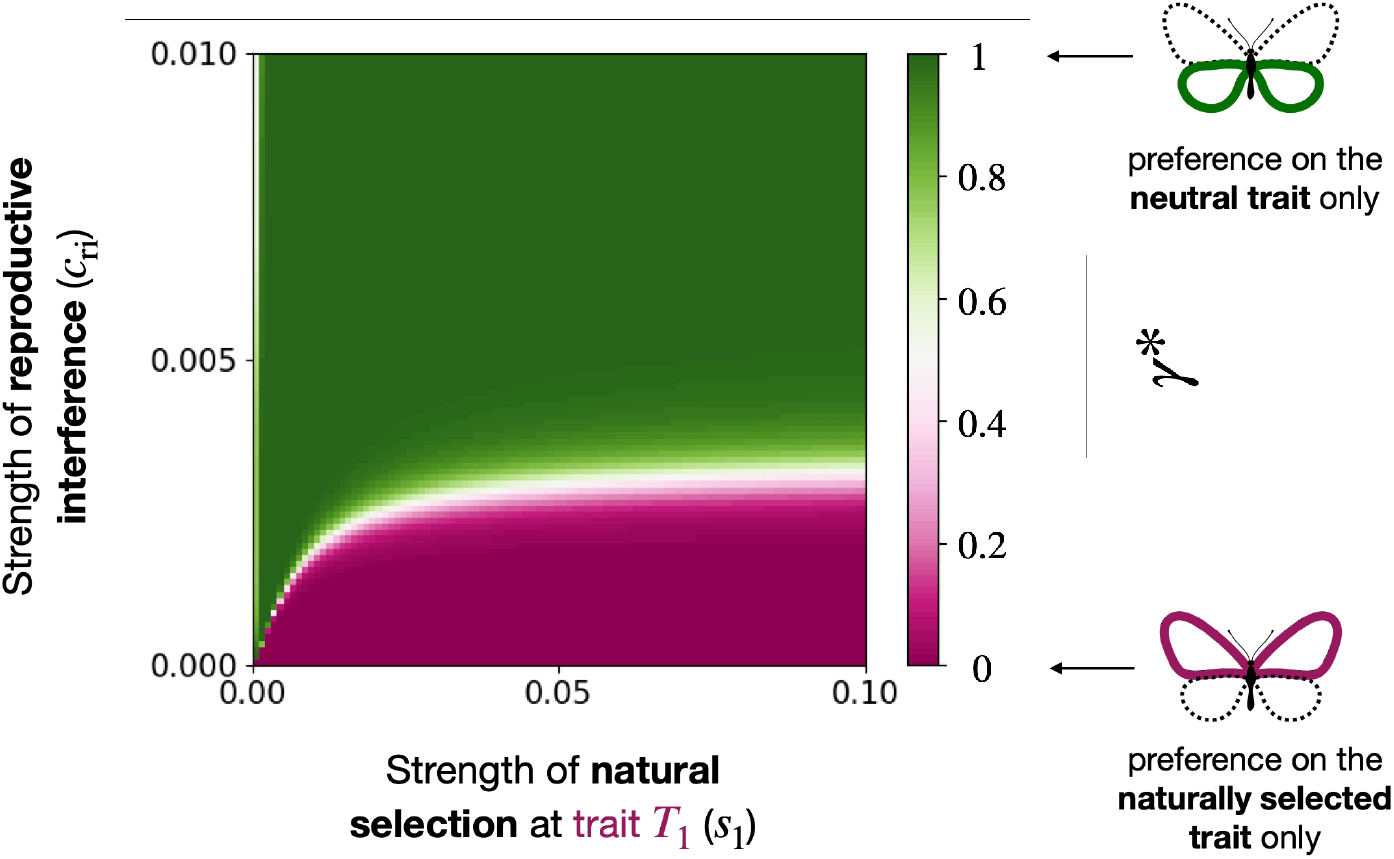
Evolution of *relative preference weighting* towards a trait under selection *T*_1_ **or a neutral trait** *T*_2_ (*γ*^∗^**), depending on the strength of natural selection acting on trait** *T*_1_ (*s*_1_**) and the strength of reproductive interference (***c*_ri_**) when** *T*_2_ **is neutral (***s*_2_ = 0).

#### The neutral trait becomes a reliable cue for species recognition

Assuming that the two species do not sexually interact (*c*_ri_ = 0), the model predicts that female preference will target the trait under selection *T*_1_ (Figure 3). In contrast, assuming reproductive interference between the two sympatric species promotes the evolution of preference targeting the neutral trait *T*_2_. A female of species *A* prefers trait value 0 (Figure A2), reducing sexual interaction with males from species *B* always displaying the trait value 1 (see Equation (A6)). Sexual selection thus increases the phenotypic divergence with heterospecific (see Equation (A1)), making the neutral trait *T*_2_ a more reliable cue for species recognition in species *A*.

Figure 4(c) confirms that the selective advantage owing to reproductive interference rises when the resident preference tends to target the neutral trait. When preference towards the neutral trait *T*_2_ increases, the stronger sexual selection enhances the phenotypic divergence with heterospecific (Figure A3), making *T*_2_ a more reliable cue for species recognition. Then in an adaptive dynamic framework, during the recurrent fixation of mutant alleles, a positive feedback loop promotes preference towards the neutral trait *T*_2_.

**Figure 4.**
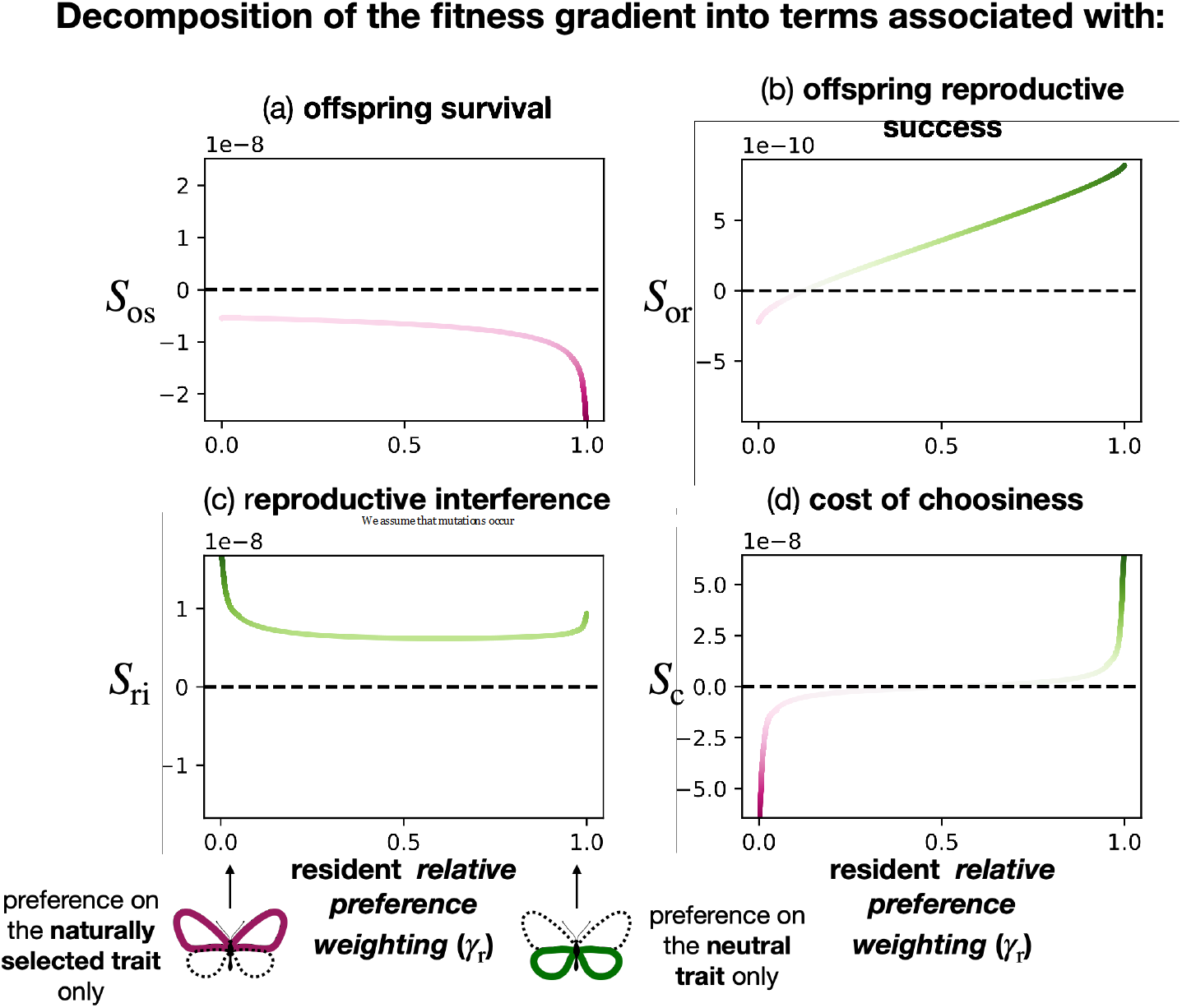
Decomposition of the fitness gradient into terms associated with (a) offspring survival (*S*_os_), (b) offspring reproductive success (*S*_or_), (c) reproductive interference (*S*_ri_) and (d) cost of choosiness (*S*_c_), depending on the resident *relative preference weighting* value (*γ*_r_). We fixed the level of reproductive interference (*c*_ri_ = 0.0025) and assume that trait *T*_1_ is under natural selection (*s*_1_ = 0.02) while trait *T*_2_ is neutral (*s*_2_ = 0). We assume cost of choosiness (*c* = 0.001). When the line is green (resp. purple), the corresponding evolutionary force (offspring survival, offspring reproductive success, reproductive interference and cost of choosiness) promotes the evolution of preference towards the neutral selection *T*_2_ (resp. the trait under selection *T*_1_). The more intense the colour, the more intense the selection.

#### Species interaction limits preference enhancing offspring fitness

Figure 3 also indicates that natural selection promotes preference towards the trait under selection *T*_1_. As long as natural selection is strong relative to reproductive interference, females from species *A* prefer males with the allele 1 at the trait under selection (Figure A2), also displayed by the males of the sympatric species *B*. However, when reproductive interference is strong relative to natural selection, females preference target the neutral trait *T*_2_ reducing heterospecific mating than enhancing offspring fitness.

Figure 4(a) reveals that the selective benefit of producing adapted offspring decreases when resident preference targets the trait under selection. When preference towards the trait under selection *T*_1_ increases, the stronger sexual selection reduces the genetic diversity (Figure A3). As most males displayed the well-adapted trait value 1 (Figure A3), the advantage of preference towards the trait under selection decreases. By contrast with the evolution of the preference towards the neutral trait, a negative feedback loop limits the evolution of preference towards the trait under selection.

Moreover, the selective benefit of producing sexy sons promotes preference towards the neutral trait (Figures 4(b) and A4). Because natural selection strongly reduces the phenotypic diversity at trait *T*_1_, almost all males display the sexy trait value at *T*_1_ (Figure A3). By contrast, males exhibiting the trait value 0 at the neutral trait *T*_2_ benefit from a sexual selection advantage compared to other males. Female preference towards males exhibiting this trait values at trait *T*_2_ is then further advantaged through an indirect benefit gained by their sexier sons (see Equation (A19)). The ‘sexy son’ advantage promotes the neutral trait *T*_2_ if the strength of reproductive interference is sufficiently high and promotes preference for the trait value 0 at the neutral trait *T*_2_ (Figure A4). This enhanced ‘sexy son’ advantage associated with female preferences toward neutral compared to adaptive traits can explain why preference toward cues reducing heterospecific mating can preferentially evolve.

Figure 3 suggests that reproductive interference should be weak for preference targeting the trait under selection *T*_1_ to evolve. However, our analyses rely on the hypothesis that all evolutionary forces are weak. Supplementary analyses, relaxing the weak selection hypothesis, show that preference targeting the trait under selection *T*_1_ can still evolve for larger values of reproductive interference, pending strong natural selection relative to reproductive interference (Figure A5).

### Evolution of preference for multiple traits enhancing offspring fitness and reducing heterospecific mating

Nevertheless, the contrasted selective pressures may promote the evolution of preference towards both traits, therefore jointly enhancing offspring fitness and reducing heterospecific mating. We thus test which values of cost of choosiness (*c*) and shapes of the trade-off function *f* (through the parameter *a*) allow the evolution of preference for multiple traits.

#### Without species interactions, the cost of choosiness promotes ancestral preference

Assuming no interaction between species (*c*_ri_ = 0), the model predicts that preference will almost always target the trait under selection *T*_1_ (Figure A6). Nevertheless, when the ancestral preference targets the neutral trait *T*_2_ and assuming a low sensory trade-off, the cost of choosiness can maintain preference toward the neutral trait *T*_2_ (Figure A6(b)).

#### Ancestral single trait preference can limit the evolution of preference for multiple traits

We then investigate the effect of ancestral preferences on the evolution of *relative preference weighting* expressed in females. We compare ancestral preferences targeting: equally both traits 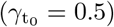, mainly the trait under selection or the neutral trait 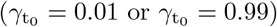.

Figures 5(a), 5(b) and 5(c) show that preferences are biased towards ancestral preference. Interestingly, ancestral preference influences the evolution of mate preference when substantial trade-offs and cost of choosiness are assumed. Under a strong trade-off, preference based on both traits effectively leads to poor attention towards both traits. This lack of attention towards both cues, therefore, creates a fitness valley preventing the switch of female attention from one trait to the alternative one. When female choice is mainly based on one trait ancestrally, positive selection promoting choice for the alternative trait may not be powerful enough to cross this fitness valley.

**Figure 5.**
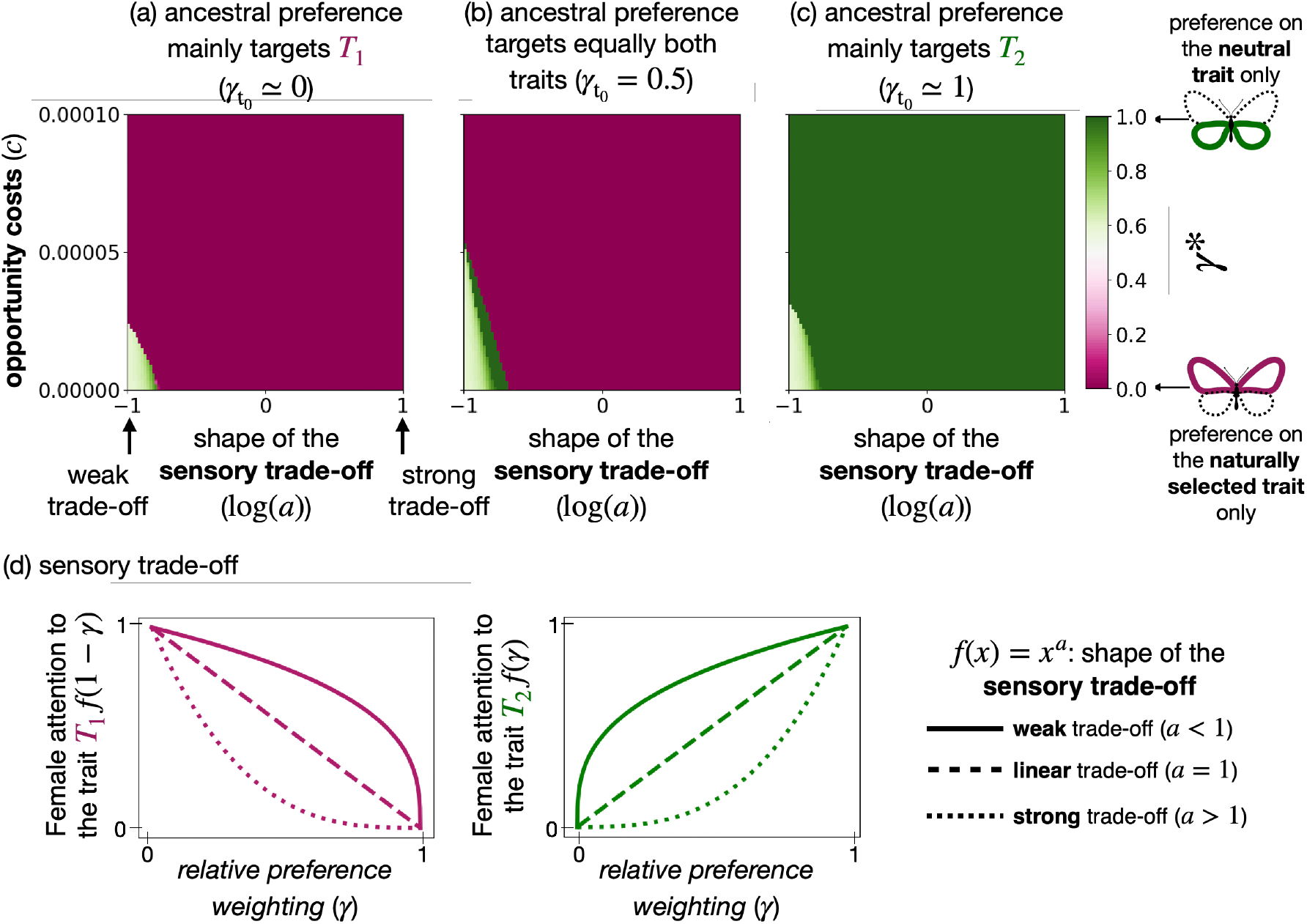
Evolution of *relative preference weighting* towards a trait under selection *T*_1_ or a neutral trait *T*_2_ (*γ*^∗^) depending on the shape of the cognitive trade-off function (through the parameter *a*) and the cost of choosiness *c* for different ancestral preferences. We assume ancestral preference targeting: (a) mainly trait *T*_1_ 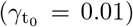, (b) equally both traits 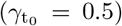, (c) mainly trait *T*_2_ 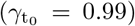. We assume reproductive interference (*c*_ri_ = 0.0025), that trait *T*_1_ is under natural selection (*s*_1_ = 0.02) and trait *T*_2_ is neutral (*s*_2_ = 0).

Figure 4(d) indicates that the cost of choosiness also favours the evolution of preference towards the ancestrally most preferred trait because this trait is then likely to be highly prevalent in the males suitable for reproduction. Mate choices mainly target the already preferred trait and then benefit from a reduced cost of choosiness (see Equation (A17)).

Moreover, when the ancestral preference mostly targets one trait, Figure 4(b) indicates that sexual selection promotes preference for this trait. When a trait is already sexy, females carrying a mutant allele increasing their attention on this trait are likely to produce more sexy sons.

These mechanisms explain why the evolution of preference strongly depends on the ancestral traits, as observed in Figure 5. Interestingly, Figure 5(b) also shows that when the ancestral preference equally targets both traits, preference targeting the trait under selection is favoured. However, this may depend on the strength of the sensory trade-off and the cost of choosiness.

#### Low cost of choosiness and weak sensory trade-off allow the evolution of preference for multiple traits

Figure 5(a) shows that a weak cognitive trade-off (log(*a*) *<* 0) and low cost of choosiness (*c*) allow the evolution of preference for multiple traits. This preference enhances offspring fitness and reduces heterospecific mating by using the trait under selection and the neutral trait, respectively. More substantial cognitive trade-offs or cost of choosiness prevent the evolution of such preference for multiple traits. Interestingly, a linear trade-off (log(*a*) = 0) leads to preference based on a single trait. A concave trade-off (*a <* 1) is thus necessary for the evolution of preference for multiple traits.

#### The cost of choosiness and the sensory trade-off promote single trait preference targeting the adaptive trait

When ancestral preference equally targets both traits, a strong sensory trade-off promotes preference towards adaptive traits (Figure 5). When assuming a large sensory trade-off (*a >* 1), the ancestral preference leads to poor attention towards both traits. This poor attention toward male traits limits the previously described ‘sexy sons’ effect that promotes preference towards the neutral trait (Figure A7).

A large cost of choosiness also promotes single-trait preference targeting the trait under natural selection *T*_1_, assuming that ancestral preference equally targets both traits. Indeed natural selection acting on trait *T*_1_ reduces phenotypic diversity in species *A* and, therefore, reduces the cost of choosiness associated with preference based on the trait *T*_1_ in this species (see Equation (A17) and Figure A8).

In contrast, an intermediate cost of choosiness allows the evolution of a preference for multiple traits with biased attention towards the neutral trait *T*_2_ (Figure 5). When the cost of choosiness is not too high, the ‘sexy son’ advantage obtained through slightly biased attention toward the neutral trait balances it. However, the preference for the adaptive trait becomes more advantageous when the cost of choosiness further increases: such preference alleviates the cost of choosiness, thus favouring the single trait preference targeting the trait *T*_1_ when the cost of choosiness becomes too large.

### Connecting ‘good genes’ with reinforcement

To explore the impact of natural selection acting on mating cues shared with other sympatric species on the evolution of female preferences, we computed the equilibrium value of the *relative preference weighting γ*^∗^ for different strengths of (1) natural selection at trait *T*_1_ and *T*_2_ (*s*_1_ and *s*_2_) and of (2) levels of reproductive interference (*c*_ri_).

#### Without species interactions, the trait under stronger selection is sexy

As expected, Figure 6(a) shows that without species interactions (*c*_ri_ = 0), female preference mainly targets the trait under stronger selection. Furthermore, Figure 6(a) also highlights that in the absence of reproductive interference, preference for multiple traits is likely to emerge when both traits are under strong selection. The impact of variation in mutation rates at the different loci is described in supplementary Figure A9.

**Figure 6.**
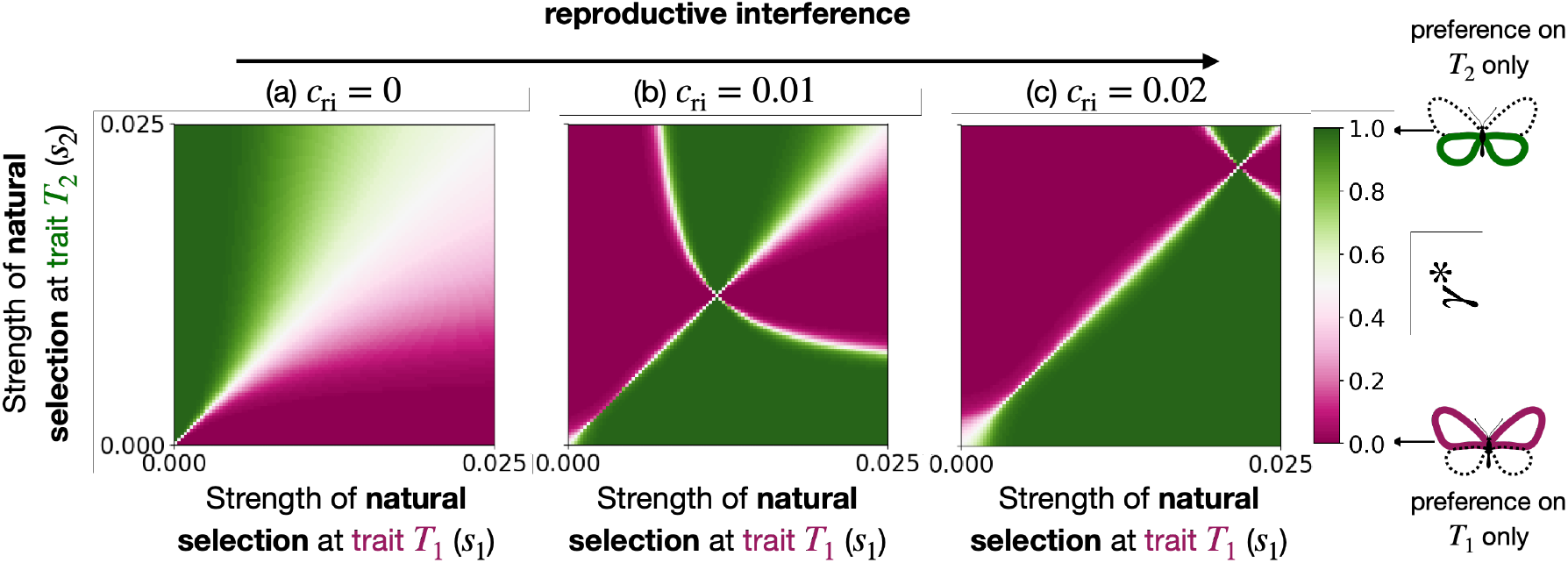
Evolution of *relative preference weighting* towards traits *T*_1_ or *T*_2_ (*γ*^∗^), depending on the strength of natural selection acting on trait *T*_1_ and *T*_2_ (*s*_1_ and *s*_2_), for different strengths of reproductive interference (*c*_ri_). We assume (a) *c*_ri_ = 0, (b) *c*_ri_ = 0.01 and (c) *c*_ri_ = 0.02.

#### Assuming reproductive interference, the trait under lower selection becomes sexy

By contrast, assuming reproductive interference with species *B* (*c*_ri_ *>* 0), females from species *A* mainly target a single trait. When natural selection on the trait *T*_1_ and *T*_2_ is low, the effect of reproductive interference prevents the evolution of preference toward the traits under the strongest selection (Figure 6(b)), because such preference increases the risk of heterospecific mating. Preference toward this trait indeed increases the frequency of the allele 0 at this locus, enhancing the reliability of this trait as cue preventing heterospecific matings, since allele 1 is fixed at both traits in species *B*. When reproductive interference further increases (Figure 6(c)), the parameter space under which a trait can become a reliable cue limiting heterospecific mating increases. Finally, Figure 6(b) and 6(c) also show that when natural selection acting on the traits *T*_1_ and *T*_2_ is very high, preference targets again the trait under the strongest selection, highlighting how natural selection may override the effect of reproductive interference.

Altogether our results show the relevance of jointly considering the effect of natural selection acting on mating cues within species, but also on the cues displayed in sympatric species to understand the evolution of mate choice.

## Discussion

Evolutionary biologists have extensively studied the evolution of mate preferences in the light of the ‘good genes’ hypothesis (Puurtinen et al., 2009) or the context of reinforcement (Servedio and Noor, 2003). By jointly considering (1) the selection regimes acting on the targeted traits within species, as well as (2) interactions with other species living in sympatry, our theoretical study provides a general framework reconciling these research fields.

We focused on natural selection regimes shared between sympatric species promoting species similarity and increasing risks of reproductive interference. For example, in the spadefoot toad, a preference for mating call increases the number of eggs fertilized in choosy females. However, it leads to reproductive interference because of the similarity of calls between sympatric species (Pfennig, 2000). Our approach drastically differs from classical studies on reinforcement, focusing on ‘magic traits’ that are under disruptive selection between species (Servedio et al., 2011). Because ‘magic traits’ are honest signals of both local adaptation and species identity, antagonistic selection regimes are not involved in the evolution of mate preferences in such a framework. Here, we investigate conflicting selections acting on the evolution of mate preferences.

We show that depending on the relative strength of natural selection and reproductive interference, females may prefer traits under natural selection, improving offspring fitness, or neutral traits, reducing heterospecific mating. Selection promotes preferences for traits under natural selection only when natural selection is strong relative to reproductive interference. Our results also show that conflicting selection may promote the evolution of preference for multiple traits. Preferences targeting multiple traits may improve offspring fitness by both transmitting adapted alleles and reducing heterospecific mating. For example, in field crickets of the genus *Teleogryllus*, females target both (1) cuticular hydrocarbons, providing fitness benefits to their offspring (Berson and Simmons, 2019), and (2) male calling song (Hill et al., 1972) that differentiates sympatric species (Moran et al., 2020).

Nevertheless, the cost of choosiness limits the evolution of such preference and can change the trait mainly targeted by female choice. The cost of choosiness promotes preference based on naturally selected traits rather than traits allowing species recognition. As natural selection erodes phenotypic diversity, preference based on traits allowing species recognition leads to stronger opportunity cost, promoting preference targeting the naturally selected traits. However, when the cost of choosiness is more limited, our model highlights that female preference may then preferentially target traits that differ from other species. For example, in *Heliconius* butterflies, where females are usually mated rapidly after emergence because of a high density of suitable males, female preference has been shown to target chemical cues differentiating sympatric species (González-Rojas et al., 2020).

Our model assumes that traits allowing limiting heterospecific mating are neutral. However, selective constraints may also act on traits used as species recognition cues. For instance, variation in such cues may influence their detection probability by predators: the conspicuousness of a trait may enhance the identification by sexual partners but may, in turn, also increase parasitism and predation risks (observed on visual, acoustic and olfactory cues in numerous organisms reviewed in Zuk and Kolluru (1998)). Increasing costs of sexual trait conspicuousness may theoretically promote the display of combinations of cryptic traits allowing recognition (Johnstone, 1995), therefore promoting preference towards multiple cues.

Our results also highlight how indirect fitness benefits and/or reproductive interference can promote female preference for multiple traits. The evolution to such a preference occurs only when the cognitive trade-off is weak. Multiple traits-based mate choice may thus preferentially evolve in species where multiple sensory systems allow such cognitive integration. Nevertheless, due to evolutionary trade-offs, the development of sensory systems is frequently associated with the regression of others (Barton et al., 1995; Nummela et al., 2013). Moreover, physical constraints may generate sensory trade-offs. For example, a visual system model of the surfperch reveals a trade-off in the performance between luminance and chromatic detection because of the limited numbers of the different types of cones in the eyes (Cummings, 2004). Neural integration of multiple information may also be limited, generating trade-offs in using multiple traits in decisions. In the swordtail fish *Xiphophorus pygmaeus*, females prefer a visual or an olfactory trait when experimenters expose them to the variation of only one out of the two traits in the potential mates. However, when both traits vary in potential mates, females do not express preference (Crapon de Caprona and Ryan, 1990), suggesting that sensory trade-off limits the use of multiple traits in preference.

Nevertheless, several alternative decision mechanisms may reduce cognitive trade-offs. For example, sequential/hierarchical mate preference, whereby targeted traits are processed in a hierarchical order, efficiently produces a decision, even considering many traits (Gigerenzer et al., 1999). Sequential mate preference is common (*e*.*g*. (Shine and Mason, 2001; Eddy et al., 2012; Gray, 2022)) and may allow the evolution of preference for multiple traits. Sequential mate choice may emerge because some traits are visible at long distances (such as colour or calls). In contrast, others are perceived only at short distances (such as oviposition site guarded by males or male-emitted pheromones) (*e.g*. (Candolin and Reynolds, 2001; *López and Martín, 2001; Mérot et al*., *2015))*.

The distance at which choosers perceive different traits may play a key role in reproductive isolation (Moran et al., 2020). Females using a cue for species recognition detectable at a short distance may have already spent time and energy or need to deploy substantial efforts to avoid heterospecific mating. Therefore, females may still suffer from increased costs associated with reproductive interference, even if they eventually manage to avoid mating with heterospecific males (Gröning and Hochkirch, 2008). Hence reproductive interference may promote preference targeting traits detected from far distances that efficiently reduce heterospecific interactions.

Reproductive isolation between species also depends on the niche of individuals of both species. Mating may preferentially occur between individuals sharing the same niche leading to niche-based assortative mating. Niche segregation may play a key role in the evolution of reproductive isolation. In two treefrog species, differing by their mating call (Park et al., 2013), different spatial and temporal segregations in calling and resting places during the breeding period increase reproductive isolation (Borzée et al., 2016). As well as sequential mate preference, niche segregation may efficiently participate in reproductive isolation without generating a trade-off with a preference for other traits.

Our study shows how natural and sexual selection may have a conflicting influence on the evolution of mate choice, specifically on the emergence of for multiple traits in sympatric species. Our study highlights the importance of (1) identifying the trade-off limiting attention towards different traits and (2) estimating the strength of the cost of choosiness to understand what traits are likely to be targeted by preference.

## Data Availability

There is no data associated with this study. Codes to generate the figures are available online at github.com/Ludovic-Maisonneuve/evolution of multiple traits preference.

## Acknowledgments

The authors would like to thank the ANR SUPERGENE (ANR-18-CE02-0019) for funding the PhD of LM, and the Emergence program from the Paris city council for supporting the team of VL. This work was also supported by the Chair “Modélisation Mathématique et Biodiversité” of VEOLIA-Ecole Polytechnique-MNHN-F.X. LM would like to thank Thomas Aubier for helpfull discussions. The authors would like to thank Judith Mank, Andy Gardner and the two anonymous reviewers for the relevant comments on previous versions of the article.

## Author Contribution

LM, CS, and VL conceived and designed the study. LM built and analyzed the model. LM wrote the manuscript with contributions from all authors. LM wrote the Python and Mathematica scripts.

## Conflict of interest

The authors of this preprint declare that they have no conflict of interest with the content of this article.

## Appendix

### A1 QLE analysis

#### Evolution of mating traits under natural and sexual selection

First, we explored the relative effects of natural and sexual selections on the evolution of traits in species A. Following the QLE approach, the change of allele 1 frequency at *T*_*i*_, for *i* ∈ {1, 2}, after one generation in this species is given by:

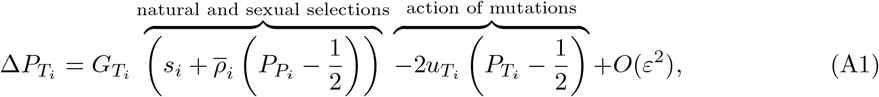

where *G*_*I*_ is the genetic diversity at locus *I* ∈ *{T*_1_, *T*_2_, *P*_1_, *P*_2_, *M}* given by

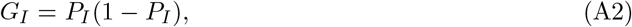

and 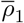 and 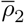are the average strengths of preference on traits *T*_1_ and *T*_2_ respectively in the population

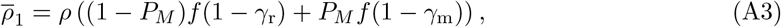

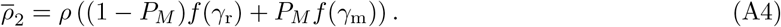

While the action of natural selection simply depends on the advantage of trait value 1 due to natural selection *s*_*i*_, the effect of sexual selection is modulated by the average strength of preference on trait *T*_*i*_ in the population 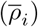. Sexual selection promotes the allele 1 when most females prefer the associated trait value 1 *i.e*. when 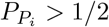.

#### Evolution of trait value preference

Now we explore the selection acting on preference loci *P*_1_ and *P*_2_, determining which trait value is sexy at traits *T*_1_ and *T*_2_. Following the QLE approach, the change of allele 1 frequency at *P*_*i*_, for *i* ∈ {1, 2}, after one generation in this species is given by:

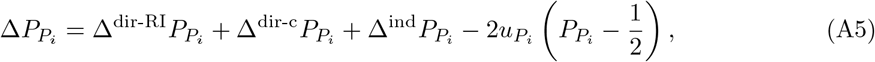

where 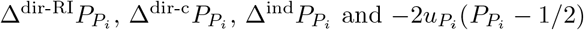 describes the effect of direct selection due to reproductive interference and cost of choosiness, indirect selection and mutations.

##### Reproductive interference promotes preference for the trait value more common within conspecific than heterospecific

The effect of reproductive interference on the change of allele 1 frequency at *P*_*i*_, for *i* ∈ {1, 2} is given by

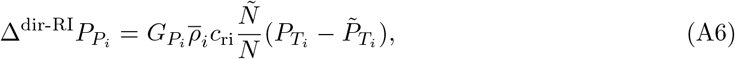

where 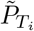 is the frequency of allele 1 at locus *T*_*i*_ in species B.

Selection acting on locus *P*_*i*_ depends on how much preference targets the trait *T*_*i*_, captured by 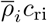. Reproductive interference promotes preference for the trait value 1 when trait value 1 is more common within conspecific than within heterospecific *i.e*. 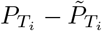
.

As expected, the effect of reproductive interference mainly depends on the density ratio between species B and A, *Ñ/N* : the probability that a female in species A encounters an heterospecific male increases with *Ñ/N*. Selection caused by reproductive interference also increases with the strength of preference *ρ*. Strong preference leads to more significant fitness differences between females, with different preferences intensifying selection due to reproductive interference.

##### Sympatry with other species intensifies the cost of choosiness

Preference allows the rejection of heterospecific males but also leads to the rejection of conspecific males. After rejecting a male, a female has a probability *c* of not encountering another male leading to an opportunity cost. The effect of these costs on the change of allele 1 frequency at *P*_*i*_, for *i* ∈ {1, 2} is given by

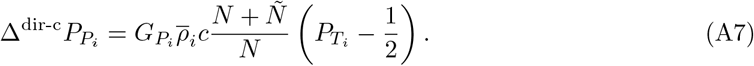

The cost of choosiness disfavours preference for trait value 1 when this trait value is the scarcest in the population *i.e*. 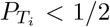. Surprisingly, selection due to cost of choosiness increases with the proportion of heterospecifics. When a female rejects a conspecific male, she must wait for another suitable male. However, females will be likely to encounter heterospecific males before encountering a conspecific male, making the rejection of a conspecific more dramatic when conspecific males are rare. The effect of the cost of choosiness is thus proportional to the average number of males a female will encounter until she encounters a conspecific (*N* + *Ñ*)*/N*.

##### Indirect selection promotes preference producing locally adapted offspring and sexy sons

Frequencies at preference loci *P*_1_ and *P*_2_ not only directly change the fitness because this modifies reproductive interference and the cost of choosiness but also because of associations with different alleles at the traits loci *T*_1_ and *T*_2_ in the offspring, leading to contrasted indirect fitness benefits. Within offspring, the preference allele at locus *P*_*i*_ becomes associated with the preferred alleles at trait *T*_*i*_ for *i* ∈ {1, 2}. Under the QLE assumptions, the genetic association between alleles 1 at loci *T*_*i*_ and *P*_*i*_, for *i* ∈ {1, 2}, is given by

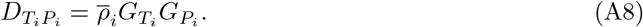

Because of mate preference, there is a positive association between the genetic basis of preference for one trait value and the genetic basis of this trait value.

The term describing the effect of indirect selection on the change of allele 1 frequency at *P*_*i*_, for *i* ∈ {1, 2} is given by

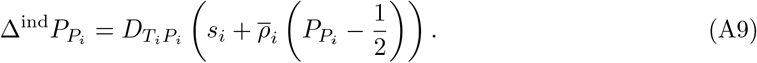

When the mutant is associated with a trait value, direct selection on this trait indirectly affects the change of mutant frequency. Then indirect selection promotes the evolution of preference towards trait value promoted by natural or sexual selection.

#### Evolution of a mutant modifying *relative preference weighting*

We investigate the evolution of the focus of female preference on either trait. We thus study the invasion of a mutant at locus *M* associated with the value *γ*_m_, differing from the value *γ*_r_ associated with the ancestral allele. Under the QLE approximation, the allele frequency variation at the preference locus can be divided into three terms, denoted ∆^dir-RI^, ∆^dir-c^ and ∆^ind^*P*_*M*_, reflecting the effect of direct selection due to reproductive interference and the cost of choosiness and indirect selection, on the change of the mutant frequency ∆*P*_*M*_ respectively.

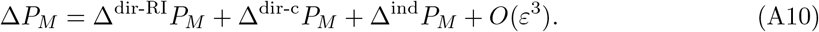

##### Reproductive interference promotes preference targeting the trait leading to strongest heterospecific avoidance

The effect of reproductive interference on the change of mutant frequency is given by

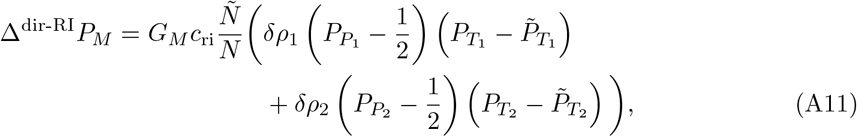

where *δρ*_1_ and *δρ*_2_ quantify the effect of the mutant allele on the preference for trait *T*_1_ and *T*_2_, respectively, compared to the resident allele:

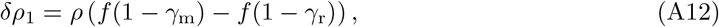

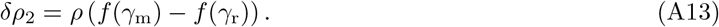

For instance, when *δρ*_2_ *>* 0, the mutant allele leads to more attention on trait *T*_2_ than the resident allele. Note that *f* is an increasing function: *δρ*_1_ and *δρ*_2_ thus have opposite signs, *i.e*. when mutant allele increases female attention on one trait, it also decreases female attention on the other trait.

Reproductive interference promotes preference for the trait allowing more accurate species recognition. Selection due to reproductive interference depends on relative phenotypic frequencies in both species. Preference for a trait leads to increased intraspecific matings than expected under random mating when the targeted trait is more common within species A than within species B. The higher the difference in trait frequencies between species, the more substantial species recognition is.

Because we have

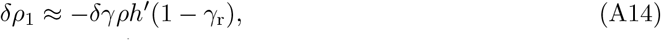

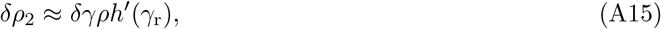

the term *S*_ri_ describing the effect of reproductive interference on mutant fitness is given by

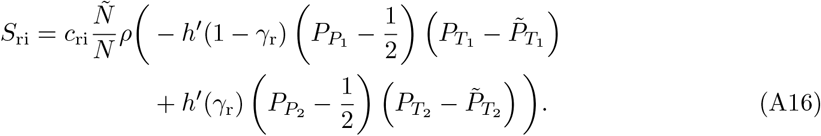

##### The cost of choosiness affects mutant fate

The effect of the cost of choosiness on mutant frequency change is given by

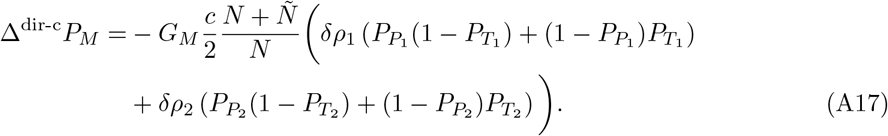

The fate of a mutant depends on the match at each trait between the most preferred trait value and the most common trait value.

The term *S*_c_ describing the effect of cost of choosiness on mutant fitness is thus given by

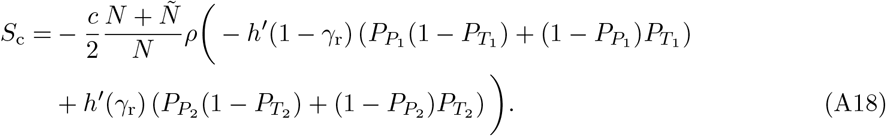

##### Indirect selection promotes preference on the trait providing the most substantial indirect fitness benefit

The term describing the effect of indirect selection on mutant alleles at locus *M* is given by

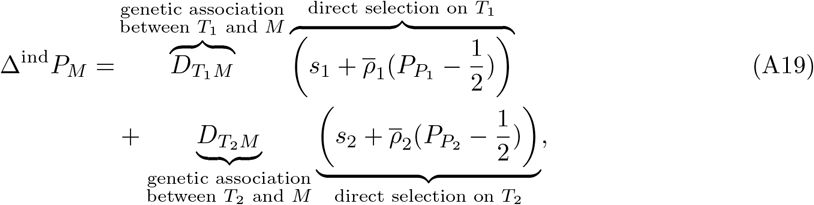

where 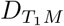 (resp. 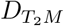) is the genetic association between the mutant allele at locus *M* and allele 1 at locus *T*_1_ (resp. *T*_2_), see (A20). When the mutant is associated with a trait value, direct selection on this trait indirectly affects the change of mutant frequency.

The genetic association between the mutant at locus *M* and the trait *T*_*i*_, for *i* ∈ {1, 2}, is given by

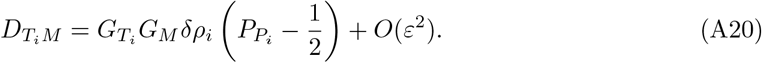

When the mutant leads to more attention on *T*_*i*_ (*δρ*_*i*_ *>* 0), the mutant becomes associated with the allele with the most preferred trait value at *T*_*i*_. Accordingly, when the mutant leads to less attention on *T*_*i*_ (*δρ*_*i*_ *<* 0), it is associated with the allele with the least preferred trait value at *T*_*i*_.

The terms *S*_os_ and *S*_or_ describing the effect of offspring survival and reproductive success on mutant fitness are thus given by

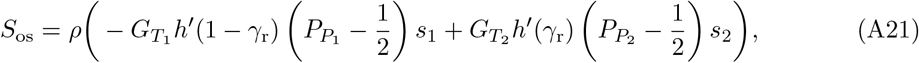

and

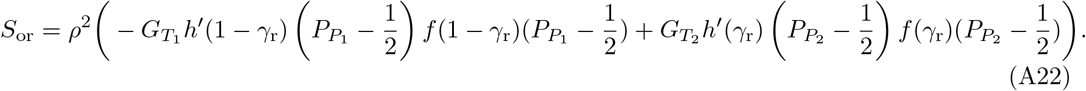

### A2 Equilibrium allele frequencies at traits and preference loci *T*_1_, *T*_2_, *P*_1_ and *P*_2_

In an adaptive dynamics framework, we study the invasion of a rare mutant at locus *M* associated with the value of *relative preference weighting γ*_m_ in a resident population where the resident allele codes for the value *γ*_r_. Before the mutant introduction, allele frequencies at traits and preference loci *T*_1_, *T*_2_, *P*_1_ and *P*_2_ evolve toward equilibrium allelic frequencies values named 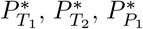 and 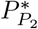. At these equilibrium frequencies, we have

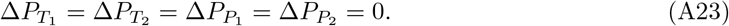

Note we may also have (A23) for different frequencies, so the reached allelic frequencies at loci *T*_1_, *T*_2_, *P*_1_ and *P*_2_ (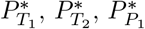and 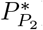) may depend on the initial frequencies at these loci.

The changes of allele frequencies at loci *T*_1_ and *P*_1_ (resp. at loci *T*_2_ and *P*_2_) do not depend on the allele frequency at loci *T*_2_ and *P*_2_ (resp. at loci *T*_1_ and *P*_1_) (see Equations (A1) and (A5)). So that before the mutant introduction, allele frequencies at loci *T*_1_ and *P*_1_ coevolve independently of allele frequencies at loci *T*_2_ and *P*_2_. We first explain how to get the value of 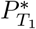 and 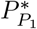 without focusing on loci *T*_2_ and *P*_2_. 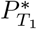 and 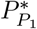 depend on the coevolution of trait and preference, only happening when resident females weight preference on trait *T*_1_. We then discriminate two cases depending on whether or not females are weighting preference on trait *T*_1_.

**Case 1: resident females weight no preference on trait** *T*_1_ (*γ*_r_ = 1)

We first compute the equilibrium allelic frequencies 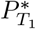 and 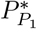 when resident females weight no preference on trait *T*_1_ (*γ*_r_ = 1). In this particular case, no selection acts at the locus *P*_1_. Because of the effect of symmetric mutations (see Equation (A5)) we have

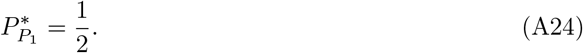

We now investigate the equilibrium allele frequencies at locus *T*_1_. Injecting 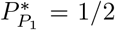 in the equation 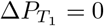 gives

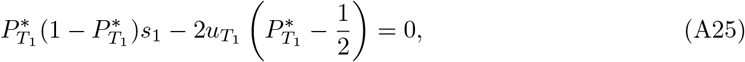

which admits a unique solution in the interval [0, 1] given by

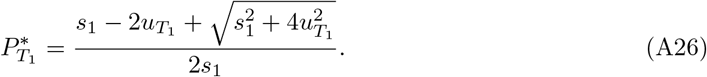

Importantly the equilibrium allele frequencies 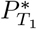 and 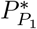 never depend on the initial allele frequency at loci *T*_1_ and *P*_1_.

**Case 2: resident females weight preference on trait** *T*_1_ (*γ*_r_ *<* 1)

We now compute the equilibrium allelic frequencies 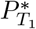 and 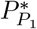 when resident females weight preference on trait *T*_1_ (*γ*_r_ *<* 1). To find candidate value of 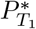 and 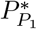 we resolve the system of equations 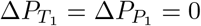. According to the equation 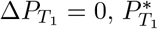 and 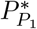 verify

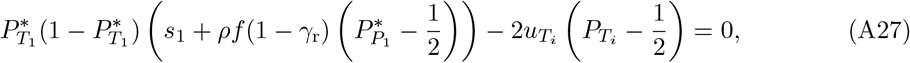

which gives

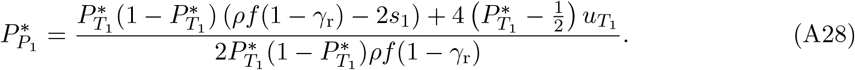

Note 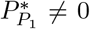 and 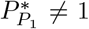 because 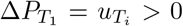 when 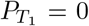and 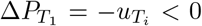 when 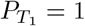 (see Equation (A1)). This ensures that Equation (A28) is always well defined.

Injecting (A28) into (A5) we find that 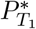 verifies (see Mathematica file in the GitHub repository)

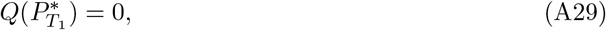

where *Q* is a polynomial function of degree 5, whose exact expression is provided in the Mathematica file.

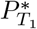 is one of the roots of the polynomial *Q*. In our model exploration, we numerically estimate the roots of *Q* within [0, 1] giving all the candidate values for 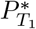. Using Equation (A28) we compute the corresponding candidate values for 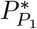. We then study the convergence stability of each pair of candidate values for 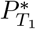 and 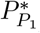. Then depending on the initial allele frequencies at loci *T*_1_ and *P*_1_, we determine the equilibrium allele frequencies reached before the mutant introduction.

We use a similar procedure to compute the values of 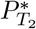 and 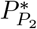.

#### Dependency on initial allele frequencies at loci *T*_1_, *T*_2_, *P*_1_ and *P*_2_

The resident population’s equilibrium allele frequencies at loci *T*_1_, *T*_2_, *P*_1_, and *P*_2_ may depend on the initial allele frequencies at these loci. Here we test this dependency for all parameter values used in our article and for all resident *relative preference weighting* values *γ*_r_. The equilibrium allele frequencies in the resident population are independent of the initial allele frequencies for all resident *relative preference weighting* values *γ*_r_ when

1. For all *γ*_r_ in [0, 1], there is a unique tuple of candidate values for 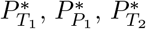 and 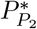.
2. This unique tuple of candidate values is convergence stable.

It appears that in the parameters space explored in this study, the equilibrium allele frequencies of the resident population are always independent of the initial allele frequencies for all resident *relative preference weighting* value *γ*_r_ (see Figure A1).

#### A3 Table and Figures

**Table A1:**
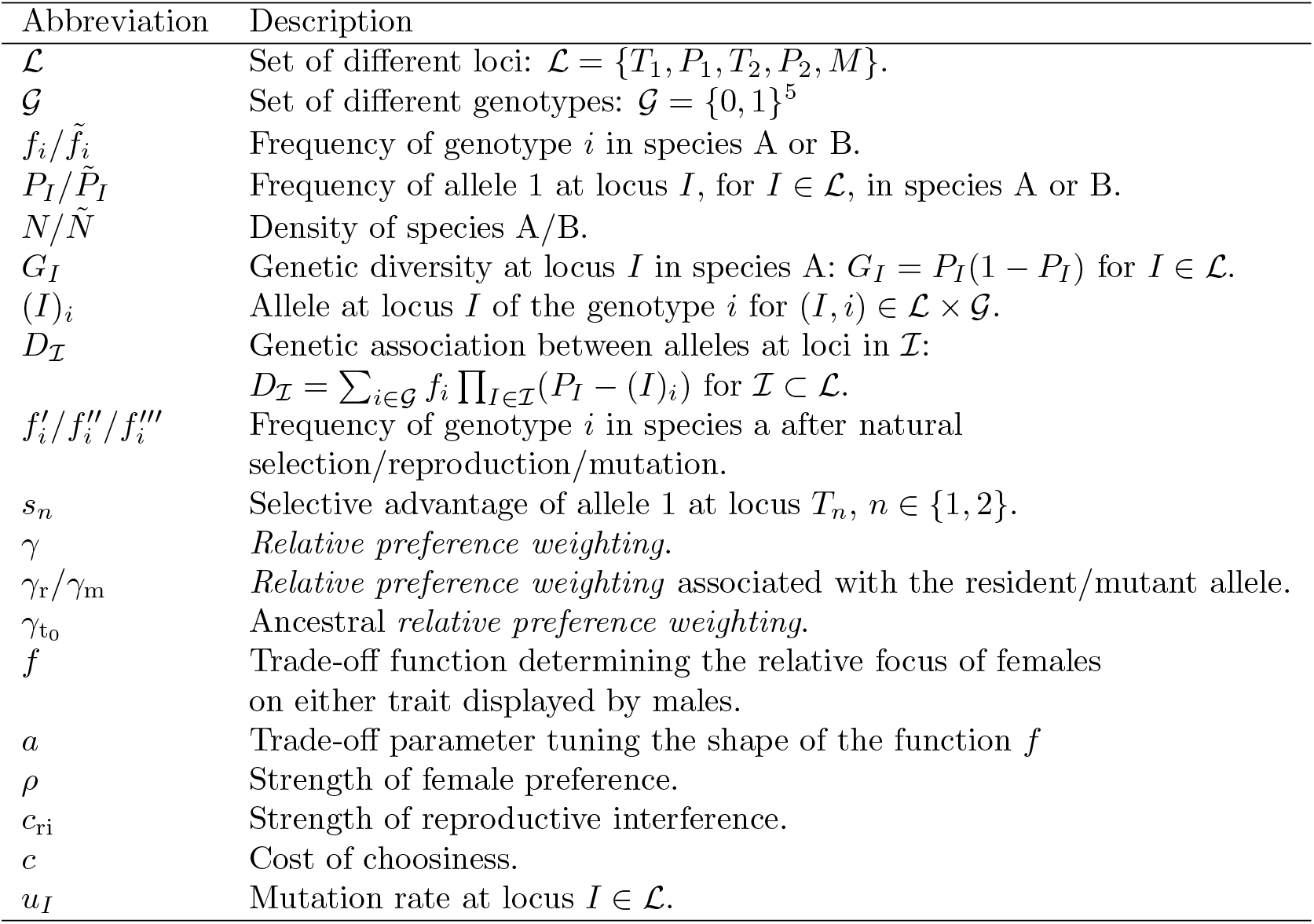
Description of variables and parameters used in the model.

| Abbreviation | Description |
| --- | --- |
| $\mathcal{L}$ | Set of different loci: $\mathcal{L} = \{T_1, P_1, T_2, P_2, M\}$. |
| $\mathcal{G}$ | Set of different genotypes: $\mathcal{G} = \{0, 1\}^5$ |
| $f_i/\tilde{f}_i$ | Frequency of genotype $i$ in species A or B. |
| $P_I/\tilde{P}_I$ | Frequency of allele 1 at locus $I$, for $I \in \mathcal{L}$, in species A or B. |
| $N/\tilde{N}$ | Density of species A/B. |
| $G_I$ | Genetic diversity at locus $I$ in species A: $G_I = P_I(1 - P_I)$ for $I \in \mathcal{L}$. |
| $(I)_i$ | Allele at locus $I$ of the genotype $i$ for $(I, i) \in \mathcal{L} \times \mathcal{G}$. |
| $D_{\mathcal{I}}$ | Genetic association between alleles at loci in $\mathcal{I}$:<br>$D_{\mathcal{I}} = \sum_{i \in \mathcal{G}} f_i \prod_{I \in \mathcal{I}} (P_I - (I)_i)$ for $\mathcal{I} \subset \mathcal{L}$. |
| $f'_i/f''_i/f'''_i$ | Frequency of genotype $i$ in species a after natural selection/reproduction/mutation. |
| $s_n$ | Selective advantage of allele 1 at locus $T_n$, $n \in \{1, 2\}$. |
| $\gamma$ | *Relative preference weighting.* |
| $\gamma_r/\gamma_m$ | *Relative preference weighting* associated with the resident/mutant allele. |
| $\gamma_{t_0}$ | *Ancestral relative preference weighting.* |
| $f$ | Trade-off function determining the relative focus of females on either trait displayed by males. |
| $a$ | Trade-off parameter tuning the shape of the function $f$ |
| $\rho$ | Strength of female preference. |
| $c_{ri}$ | Strength of reproductive interference. |
| $c$ | Cost of choosiness. |
| $u_I$ | Mutation rate at locus $I \in \mathcal{L}$. |

**Figure A1:**
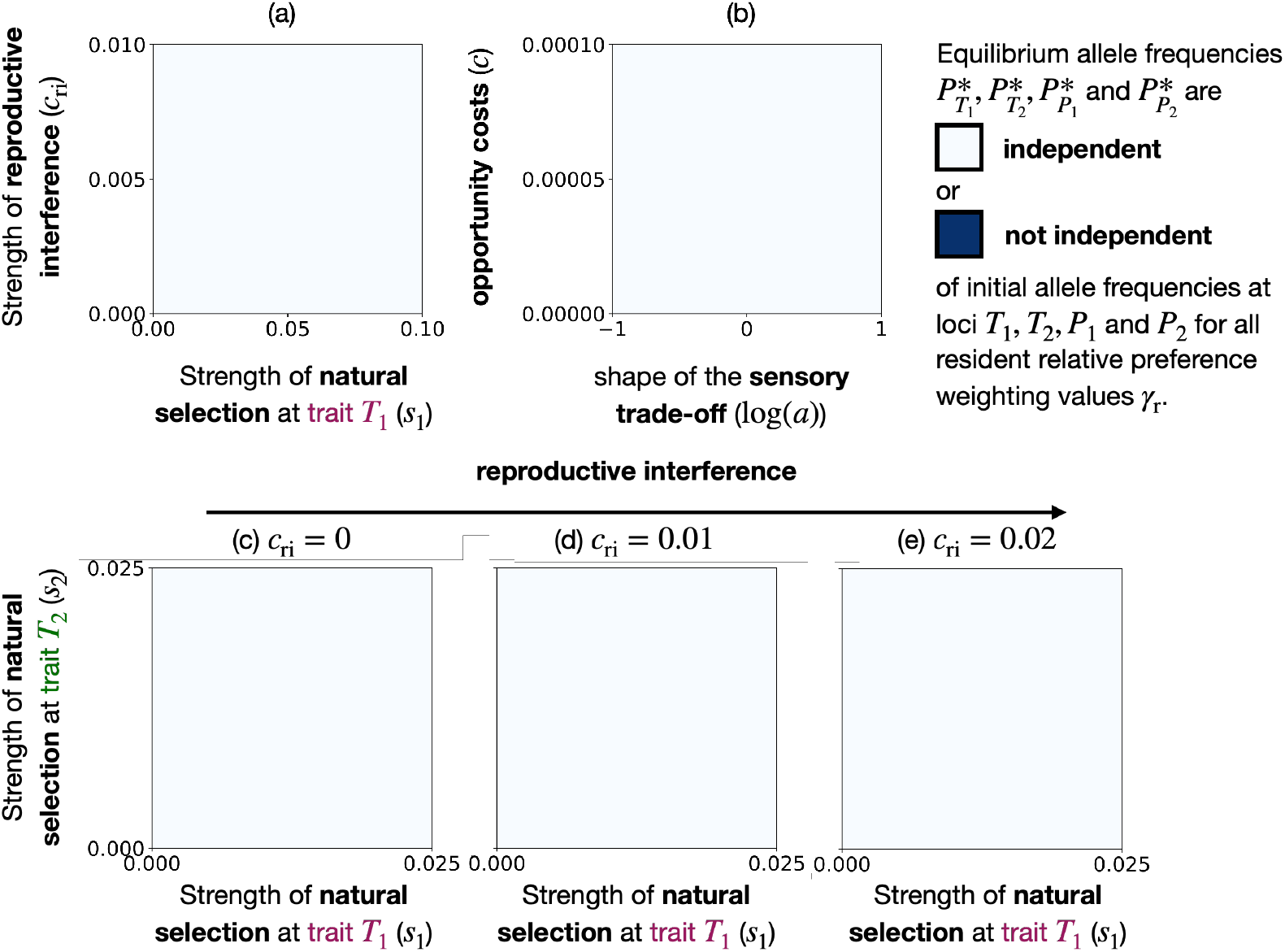
Equilibrium allele frequencies at loci *T*_1_, *T*_2_, *P*_1_ **and** *P*_2_ **in the resident population is always independent of the initial allele frequencies at these loci for all resident *relative preference weighting* values** *γ*_r_. Parameters values same as (a) Figure 3, (b) Figure 5, (c) Figure 6(a), (d) Figure 6(b) and (e) Figure 6(c).

**Figure A2:**
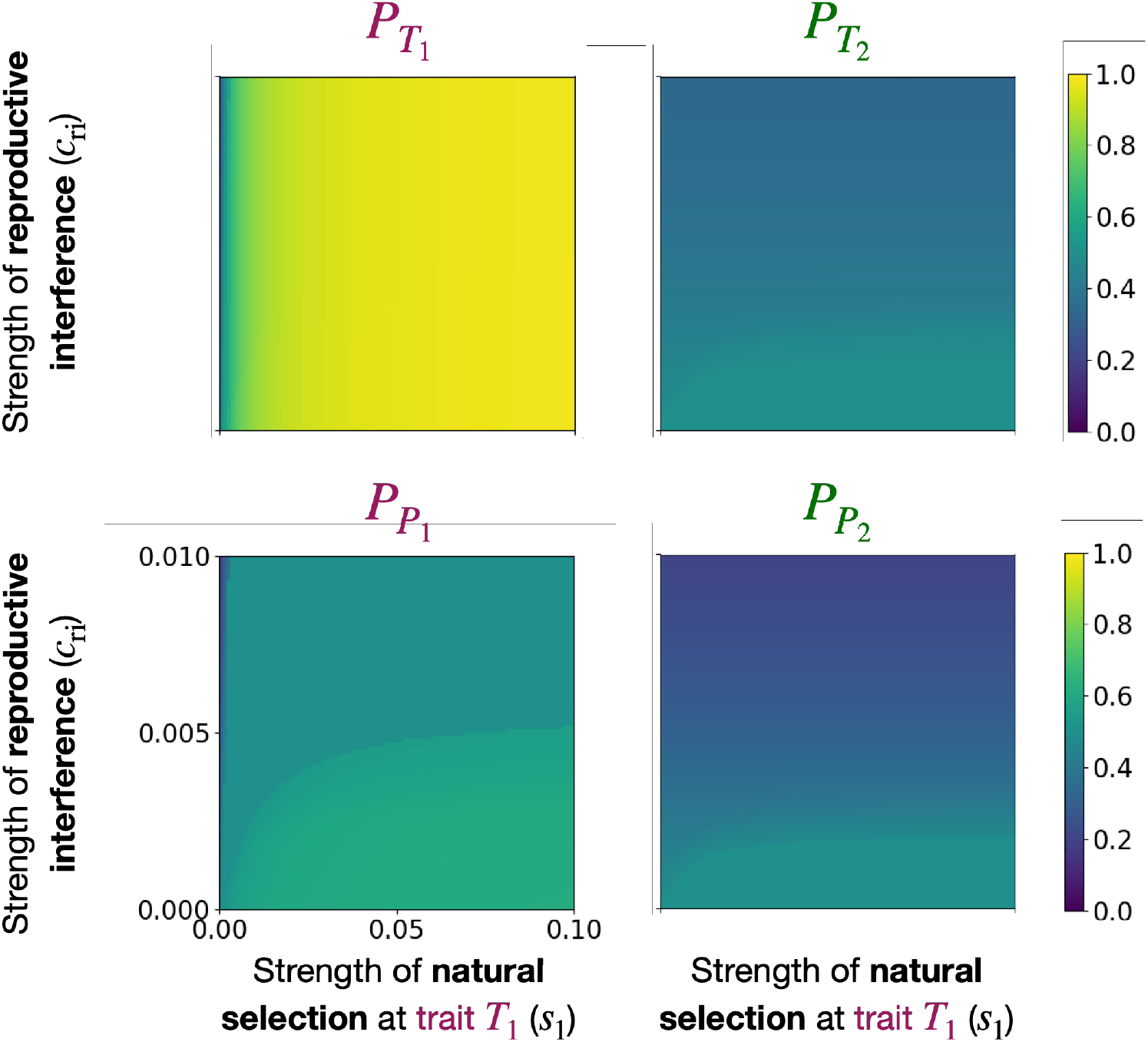
Frequency of allele 1 at loci *T*_1_, *T*_2_, *P*_1_ and *P*_2_ at equilibrium preference (*γ*_r_ = *γ*^∗^), depending on the strength of natural selection acting on trait *T*_1_ (*s*_1_) and the strength of reproductive interference (*c*_ri_), when *T*_2_ is neutral (*s*_2_ = 0).

**Figure A3:**
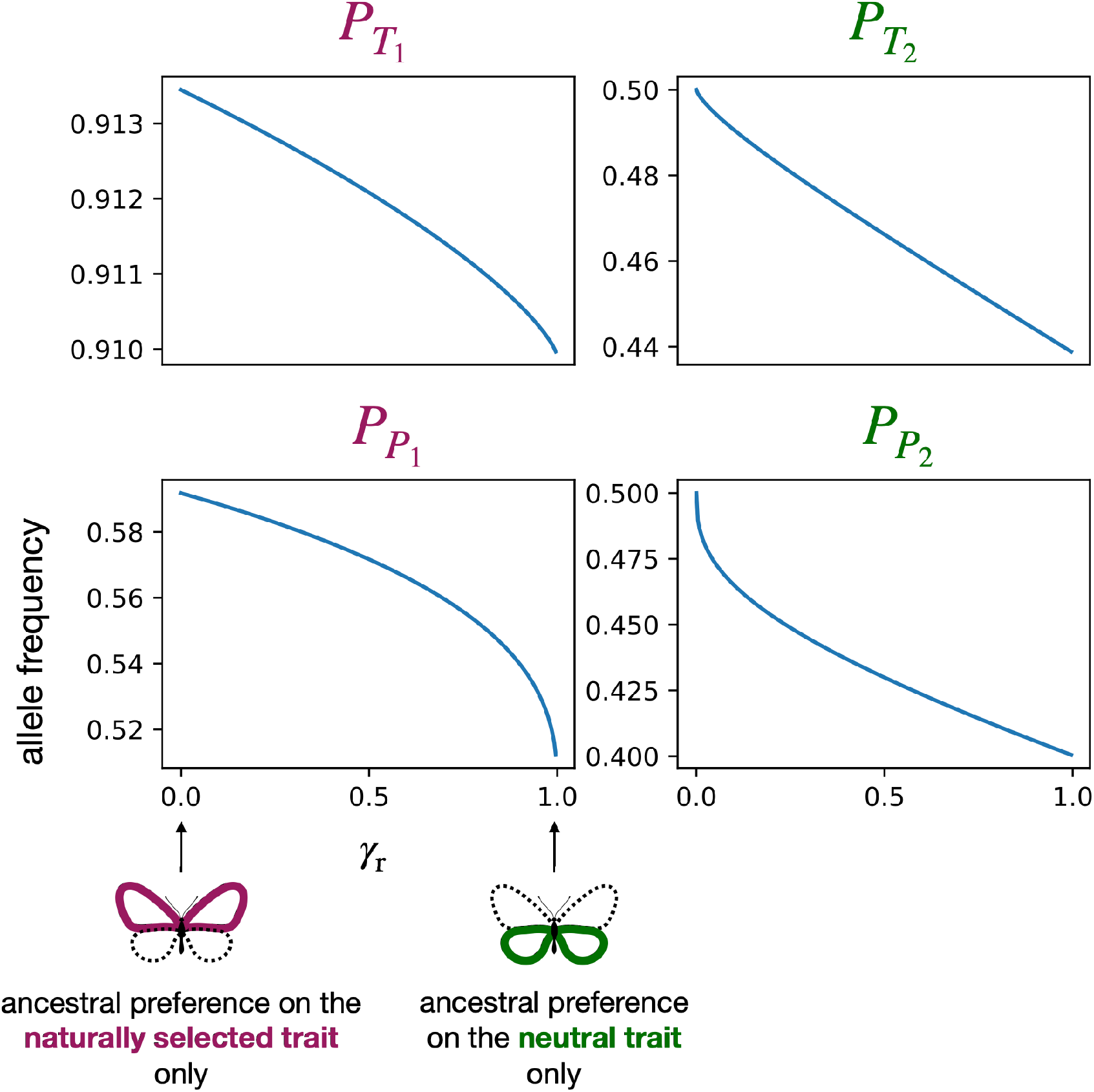
Frequency of allele 1 at loci *T*_1_, *T*_2_, *P*_1_ and *P*_2_ depending on the resident *relative preference weighting* value (*γ*_r_). We assume reproductive interference (*c*_ri_ = 0.001), that trait *T*_1_ is under natural selection (*s*_1_ = 0.05) and trait *T*_2_ is neutral (*s*_2_ = 0).

**Figure A4:**
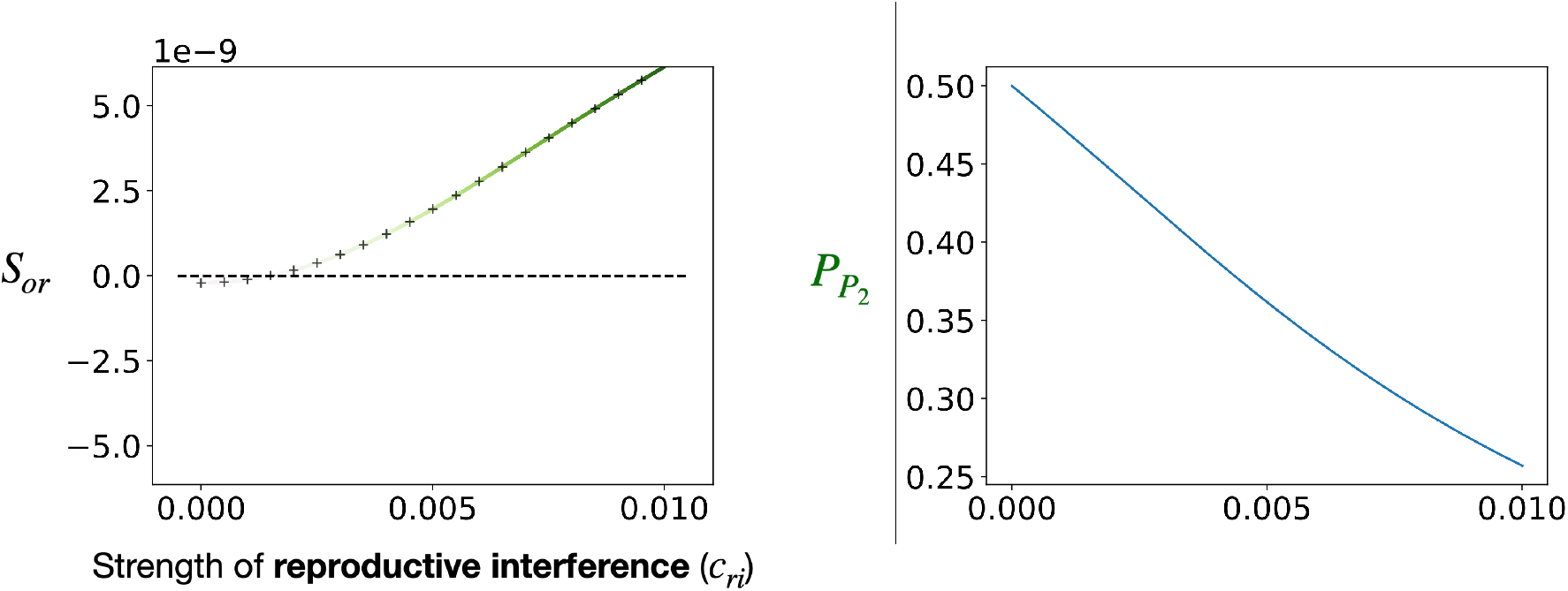
Ancestral part of the selection gradient associated with offspring reproductive success (*S*_or_) and frequency of allele 1 at locus 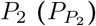 depending on the strength of reproductive interference (*c*_ri_). We assume ancestral preference targeting equally both traits 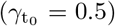. We assume reproductive interference (*c*_ri_ = 0.0025), that trait *T*_1_ is under natural selection (*s*_1_ = 0.02) and trait *T*_2_ is neutral (*s*_2_ = 0). When the line is green, offspring reproductive success promotes the evolution of preference towards the neutral selection *T*_2_. The more intense the colour, the more intense the selection.

**Figure A5:**
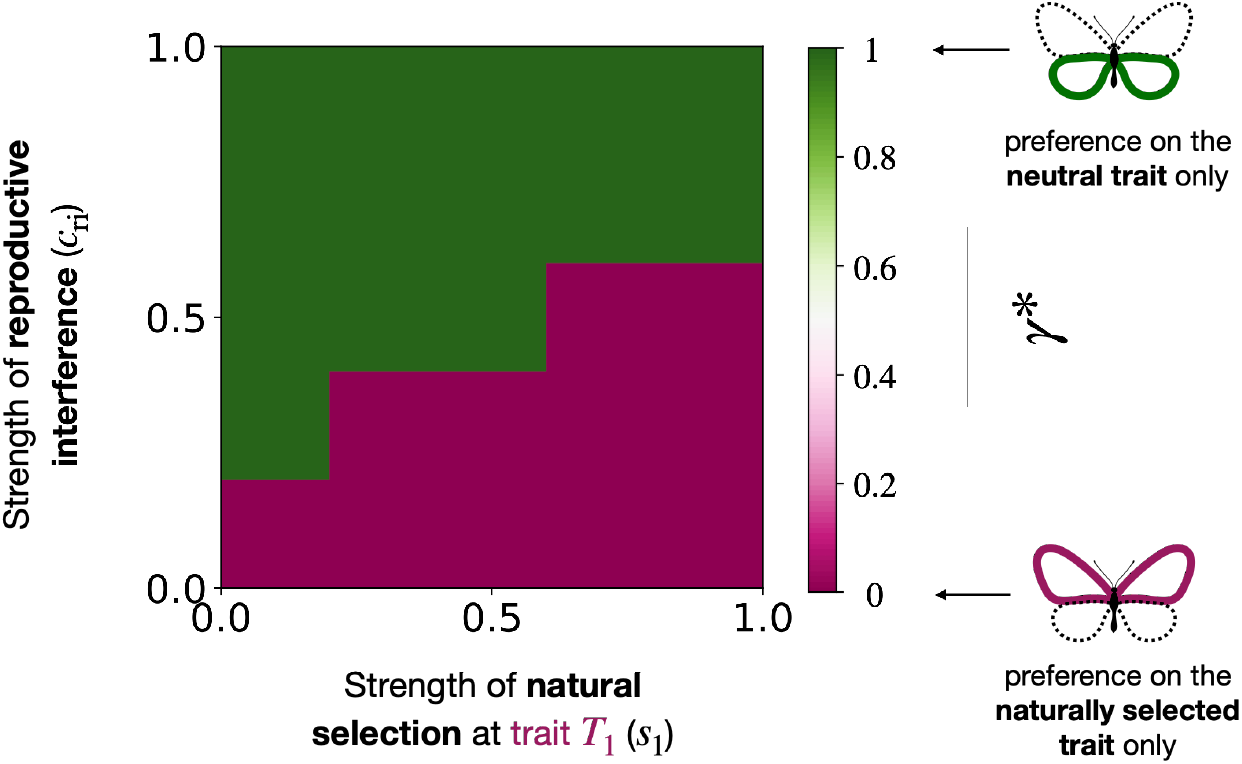
Evolution of *relative preference weighting* towards a trait under selection *T*_1_ or a neutral trait *T*_2_ (*γ*^∗^), without the QLE assumptions, depending on the strength of natural selection acting on trait *T*_1_ (*s*_1_) and the strength of reproductive interference (*c*_ri_) when *T*_2_ is neutral (*s*_2_ = 0). We assume *ρ* = 1, *c* = 0.5, 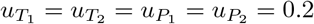.

**Figure A6:**
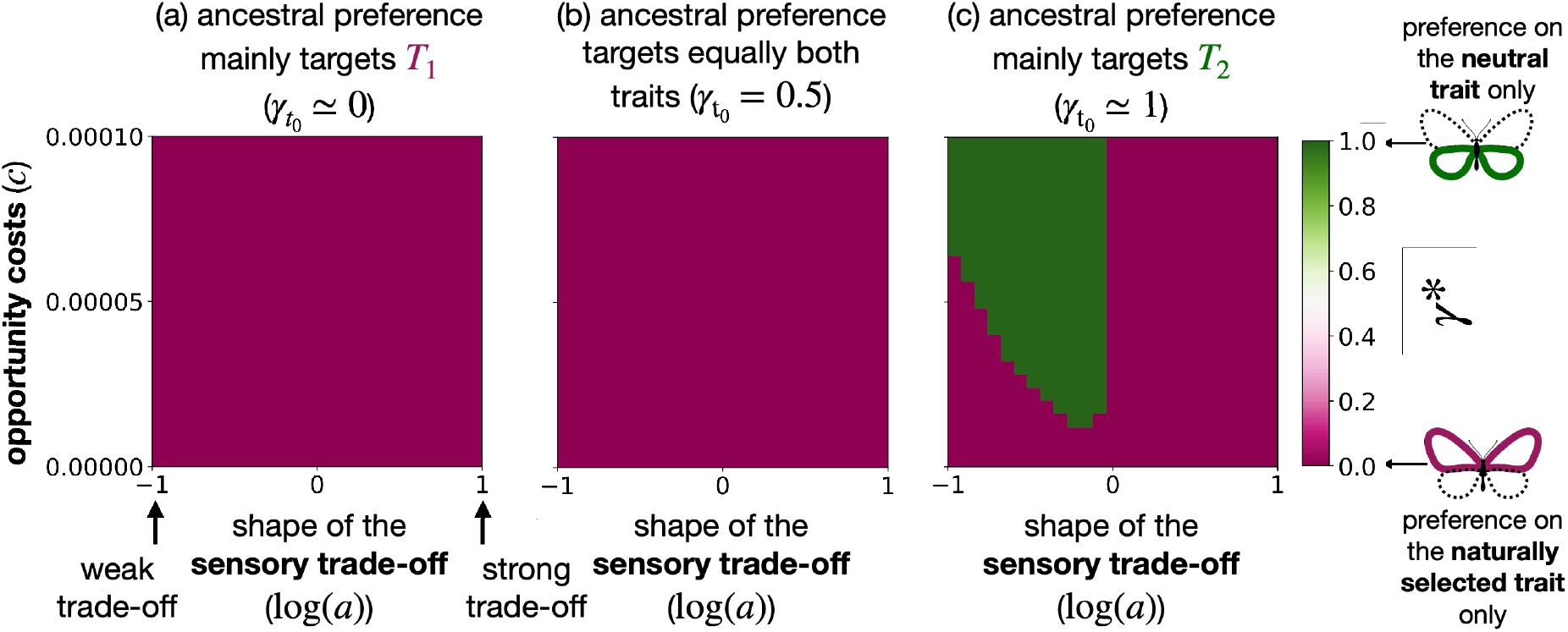
Evolution of *relative preference weighting* towards a trait under selection *T*_1_ or a neutral trait *T*_2_ (*γ*^∗^) depending on the shape of the cognitive trade-off function (through the parameter *a*) and the cost of choosiness *c* for different ancestral preferences without species interactions *c*_ri_ = 0. We assume ancestral preference targeting: (a) mainly trait 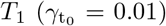, (b) equally both traits 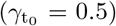, (c) mainly trait 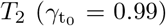. We assume that trait *T*_1_ is under natural selection (*s*_1_ = 0.02) and trait *T*_2_ is neutral (*s*_2_ = 0).

**Figure A7:**
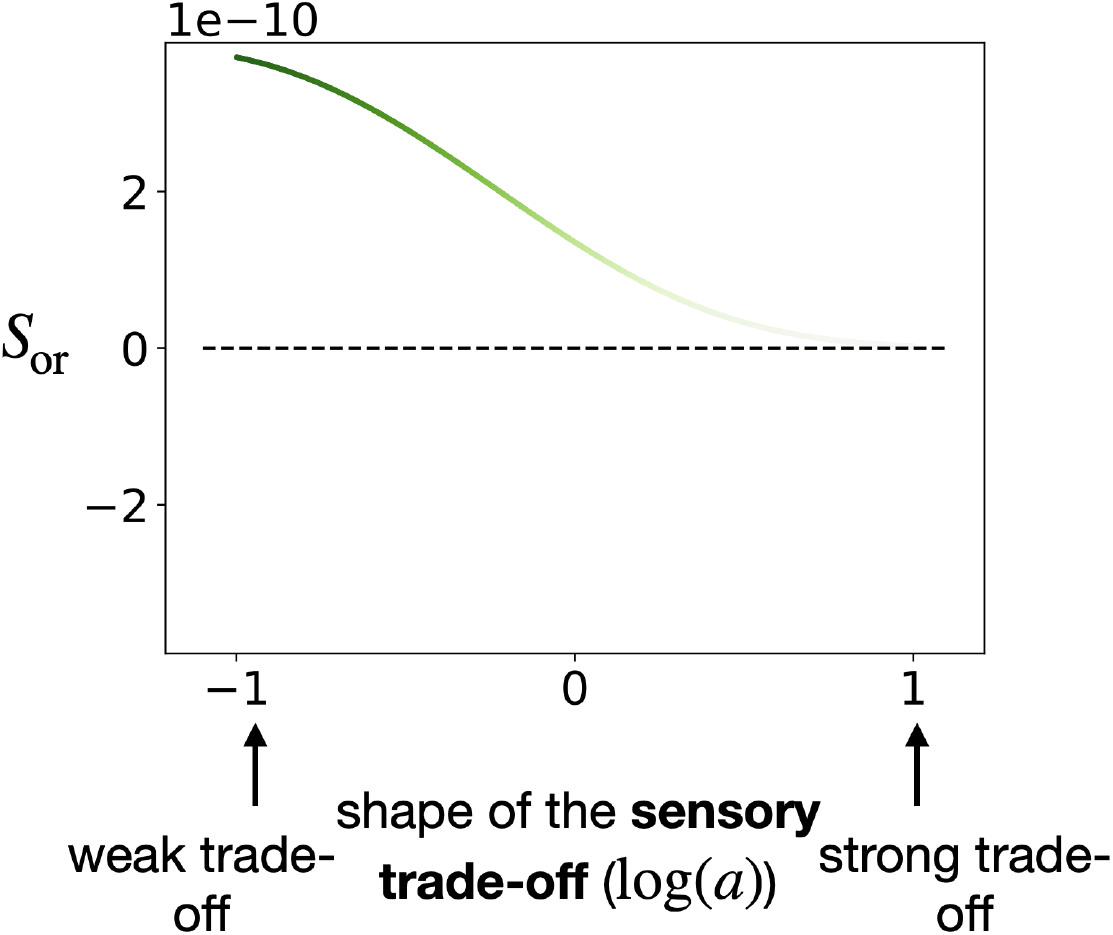
Ancestral part of the selection gradient associated with offspring reproductive success (*S*_or_) depending on the shape of the trade-off function (through the parameter *a*). We assume ancestral preference targeting equally both traits 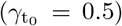. We assume reproductive interference (*c*_ri_ = 0.025), that trait *T*_1_ is under natural selection (*s*_1_ = 0.02) and trait *T*_2_ is neutral (*s*_2_ = 0). When the line is green, offspring reproductive success promotes the evolution of preference towards the neutral selection *T*_2_. The more intense the colour, the more intense the selection.

**Figure A8:**
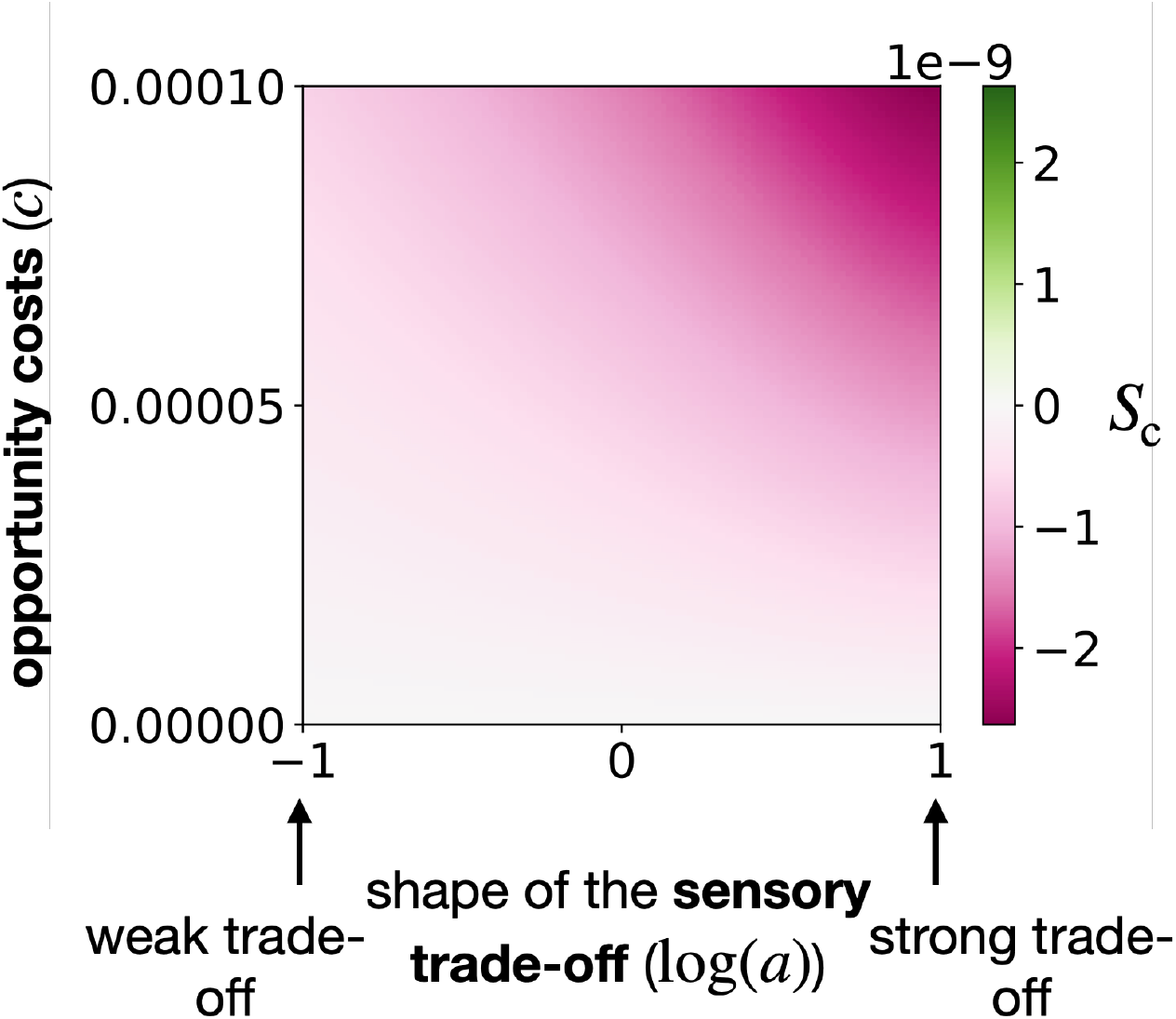
Ancestral part of the selection gradient associated with cost of choosiness (*S*_c_). We investigate different shapes of the trade-off function (through the parameter *a*) and the cost of choosiness *c*. We assume ancestral preference targeting both traits equally 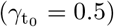. We assume reproductive interference (*c*_ri_ = 0.025), that trait *T*_1_ is under natural selection (*s*_1_ = 0.02) and trait *T*_2_ is neutral (*s*_2_ = 0). Purple area indicates that the cost of choosiness promotes the evolution of preference towards the trait under selection *T*_1_.

**Figure A9:**
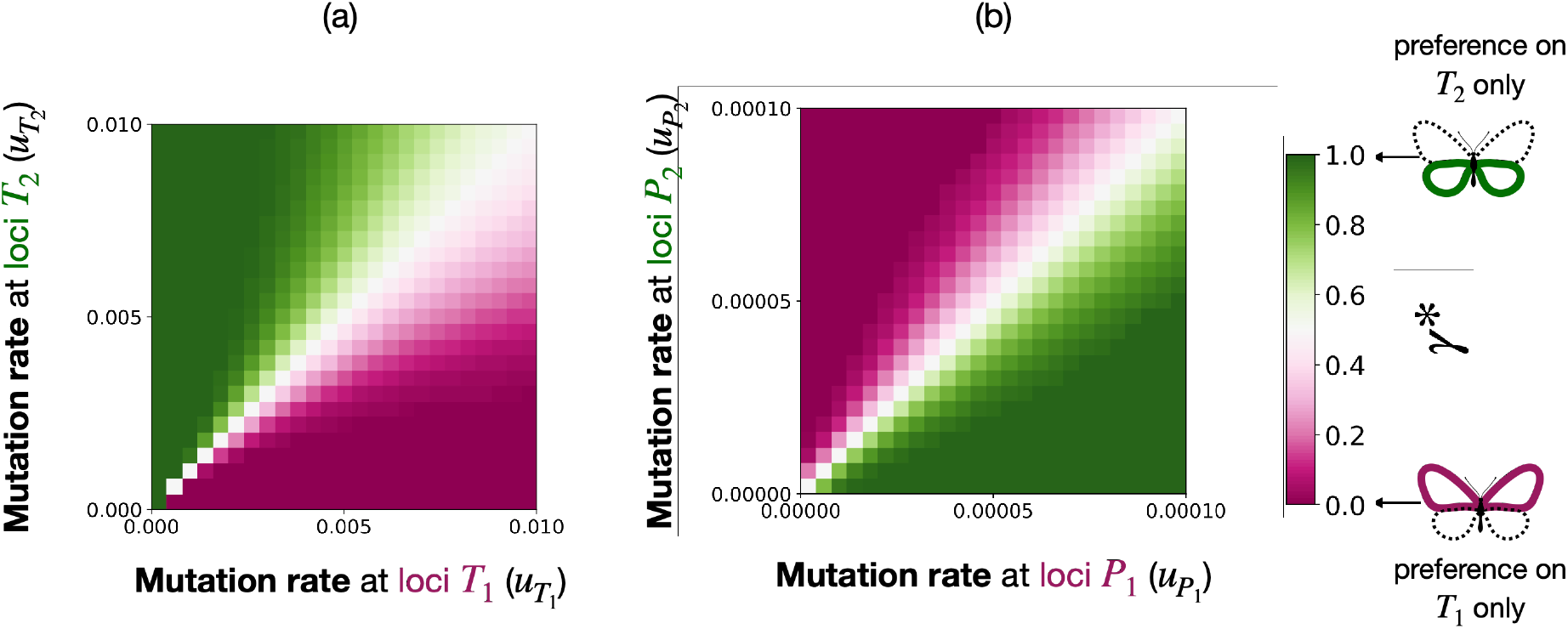
Evolution of *relative preference weighting* towards traits *T*_1_ or *T*_2_ (*γ*^∗^), depending on (a) the mutation rates at loci *T*_1_ and *T*_2_ (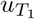 and 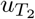) and on (b) the mutation rates at loci *P*_1_ and *P*_2_ (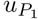 and 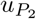), without species interaction (*c*_ri_ = 0). We assume that both traits are under natural selection (*s*_1_ = *s*_2_ = 0.02). When mutation rates at the two trait loci (*T*_1_ and *T*_2_) differ, the model predicts that female will prefer the traits associated with the highest mutation rate (see Figure A9). Mutations increase the number of males with a maladapted trait value. Such preference reduce mating with males with the maladapted trait value. When assuming that the mutation rate can differ at the loci *P*_1_ and *P*_2_, determining the preferred allele at trait *T*_1_ and *T*_2_ respectively, the model predicts that female will prefer the trait targeted by the preference locus with the lowest mutation rate (see Figure A9). Mutations at preference loci *P*_1_ and *P*_2_ indeed increase preference for the maladapted trait value, decreasing the likelihood of producing locally adapted offspring.

## Notes

### Competing Interest Statement

The authors have declared no competing interest.

### Summary of Updates

Improving the clarity and the grammar

